# ACSS2 upregulation enhances neuronal resilience to aging and tau-associated neurodegeneration

**DOI:** 10.1101/2024.03.27.586865

**Authors:** Naemeh Pourshafie, Hong Xu, Kechun Yang, Greg Donahue, Xue Lei, Shuo Zhang, Oksana Shcherbakova, Connor Hogan, Michael Gilbert, Kevt’her Hoxha, Lesley Chaboub, Virginia Lee, Peter Adams, John A. Dani, Nancy Bonini, Shelley Berger

## Abstract

Epigenetic mechanisms, including histone acetylation, are pivotal for learning and memory, with a role in neuronal function in Alzheimer’s disease and Related Dementia (ADRD). Acetyl-CoA synthetase 2 (ACSS2), an enzyme that generates acetyl-CoA, is central to histone acetylation and gene regulation, particularly in neurons, due to their unique metabolic demands and postmitotic state. ACSS2 can be recruited to the nucleus and chromatin, locally supplying acetyl-CoA to directly fuel histone acetyltransferase enzymes and key neuronal gene expression. This regulatory mechanism may be a promising target for therapeutic intervention in neurodegenerative diseases. Previously we showed that systemic ACSS2 deletion in mice, although largely normal in physiology, is greatly impaired in memory. Here we investigated whether increasing ACSS2 levels could protect neurons against disease and age-associated cognitive decline. Given the role of tau in ADRD, we used primary hippocampal neurons that mimic the sporadic development of tau pathology and the P301S transgenic mouse model for tau-induced memory decline. Our results show that ACSS2 upregulation mitigates tau-induced transcriptional alterations, enhances neuronal resilience against tau pathology, improves long-term potentiation, and ameliorates memory deficits. Expanding upon these findings, we reveal that increasing histone acetylation through ACSS2 upregulation improves age-associated memory decline. These findings indicate that increasing ACSS2 is highly effective in countering age- and tau-induced transcriptome changes, preserving elevated levels of synaptic genes, and safeguarding synaptic integrity. We thus highlight ACSS2 as a key player in the epigenetic regulation of cognitive aging and ADRD, providing a foundation for targeted therapeutics to enhance brain resilience and function.

**Summary:** ACSS2 upregulation protects neurons from disease and age-related decline by enhancing synaptic and longevity gene expression.

## INTRODUCTION

The pursuit of healthy aging through epigenetic regulators (*1–3*) is an exciting frontier in the field of neuroscience and neurodegenerative diseases. It is well-established that aging is a pivotal risk factor for many neurodegenerative diseases (*4–6*), including Alzheimer’s disease and related dementias (ADRD). Therefore, slowing down or reversing epigenetic mechanisms underpinning the aging process represents a strategic approach to mitigate the risk of brain-related diseases. Aging correlates with gradual decline in control of gene expression (*7–9*). Central to this decline is the brain which must efficiently respond to external stimuli. Brain vulnerability is particularly evident with age, with compromise of epigenetic homeostasis, which maintains and regulates gene expression, resulting in DNA hypermethylation and reduced histone acetylation (*10, 11*), among other aberrant chromatin changes. The gradual reduction of transcriptional control that occurs with age subsequently diminishes neuronal resilience, rendering the brain susceptible to decline and disease. One profound consequence of age-related decline is loss of synaptic integrity (*12–14*)—a core aspect of neuronal function. Inevitably, loss of synaptic integrity is linked to the erosion of memory and onset of dementia (*15*), thus restoration and maintenance of neuronal resilience are key strategies to counteract cognitive decline. Recently, transcription factor-based reprogramming reveals epigenetic modulation as key to aging rejuvenation (*1, 2*), underscoring the potential of epigenetic mechanisms to restore expression of genes crucial to maintain optimal neuronal function.

Central to epigenetic regulation are nuclear-localized metabolic enzymes such as Acetyl-CoA synthetase 2 (ACSS2), which catalyzes acetyl-CoA from acetate. In neurons, histone acetylation can be heavily dependent on the activity of ACSS2 (*16*). We have shown that ACSS2 resides on chromatin generating acetyl-CoA to supply key histone acetyltransferases (HATs) (*16, 17*), allowing rapid upregulation of critical neuronal genes central to learning and memory (*16, 18*). ACSS2 knockout leads to animals defective in memory; upregulation of ACSS2 has effects to boost some synaptic genes in an amyloid-beta expressing mouse (*19*). However, in contrast to amyloid-beta, tau is the toxic protein associated with memory decline in Alzheimer’s disease and other tauopathies (*20–22*), and it is unknown if ACSS2 can protect from tau or age-associated cognitive decline. In neurodegenerative disease and aging, there is a global shift to histone hypoacetylation with reduction of H4K12ac, H4K5ac, H3K14ac, and H2BK5ac in aged mice (*23*), and dramatic losses of H4K16ac, H3K18ac, and H3K23ac in human AD post-mortem brain tissue (*24, 25*). Given the impact of epigenetic modifications on gene expression, these histone acetylation losses may contribute to neural decline and degeneration (*26*).

We employed a comprehensive multidisciplinary approach to investigate whether enhancing ACSS2-dependent chromatin processes can protect neurons against age- and tau-associated epigenetic dysregulation and prevent associated cognitive decline. Our results show that upregulation of ACSS2 counteracts age-associated transcriptomic changes to maintain elevated levels of synaptic genes, thereby safeguarding synaptic integrity over time and protecting against disease-induced and age-related memory decline. Unexpectedly, upregulation of ACSS2 also prevents accumulation of pathogenic tau in vitro and in vivo, and in vivo the hippocampal neurons are rescued using electrophysiological measures. ACSS2 is a potential strategy to enhance neuronal resilience against ADRD and normal age-associated cognitive decline.

## RESULTS

### ACSS2 upregulation maintains histone acetylation and synaptic gene expression in hippocampal neurons over time

We transduced cultured mouse primary neurons isolated from the hippocampus with adeno-associated virus (AAV)-PHP.eB-ACSS2 (hereafter AAV.ACSS2). In this vector, the constitutive elongation factor 1 (EF1a) promoter drives expression of mouse ACSS2 tagged with a C-terminal flag and the mPlum fluorophore. A p2A self-cleaving peptide facilitates the separation of ACSS2 flag from mPlum, allowing the production of both proteins from the same transcript (AAV.ACSS2) (**Fig. 1A**). The viral vector expressing mPlum (“control vector”) was used as a negative control throughout the experiments. Consistent with our previous finding establishing ACSS2 as a nuclear-metabolic enzyme in neurons (*16*), we detected exogenous ACSS2 upregulation via western blotting (**Fig. S1A**), and within the nucleus of mouse hippocampal neurons by immunofluorescence (**Fig. 1B**). In accordance with the established role of ACSS2 to promote histone acetylation (*16–18*), we observed increased levels of transcription-linked H3K9ac and H3K27ac in neurons transduced with AAV.ACSS2 compared to vector control (**Fig.1, C to F**). We note that neurons exhibited clear intrinsic reduction in the level of these histone acetylation sites over time, and this trend was countered by continuous acetylation promoted by ACSS2 upregulation (**Fig.1, C to F**). We evaluated potential toxicity linked to ACSS2 upregulation via quantification of neuronal survival following viral transduction and found no reduction of neuron viability (**Fig. S1B**).

**Fig 1.**
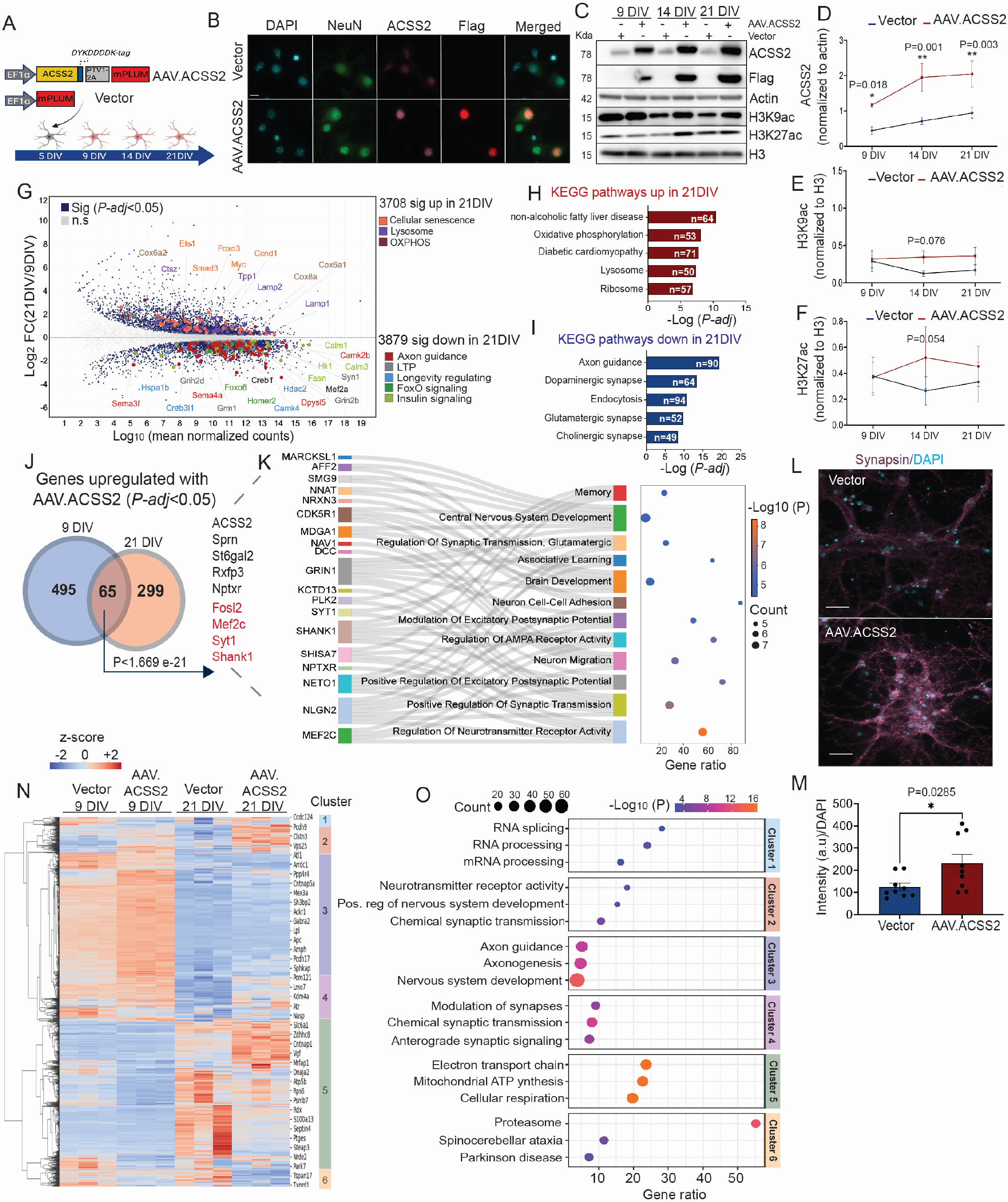
ACSS2 upregulation maintains neuronal gene expression over time. **A.** Schematic of experimental design. **B**. 60x IF images of 21 DIV hippocampal neurons (NeuN), showing nuclear localization of ACSS2 and Flag, Scale bar 50 µm. **C**. Western blot. **D**. Ratio of ACSS2 normalized to actin, **E.** H3K9ac, **F.**, and H3K27ac normalized to H3 (n=4, biologically independent samples, mean ± s.e.m, 2-way ANOVA, Fisher’s LSD test *p<0.05, **p<0.01). **G**. MA plot of RNA-seq data (n=3). Significant DEGs (*P-adj* < 0.05) are highlighted in dark blue and non-significant genes are displayed in gray. Significant genes of interest are color-coded according to their respective KEGG pathways. **H.** Top 5 KEGG pathways for upregulated and **I.** downregulated DEGs. **J.** Venn diagram showing the overlap of genes (*P-adj* < 0.05) upregulated by AAV.ACSS2/vector in 9 and 21 DIV hippocampal neurons, permutation test *P < 1.669 e-21*. Gene list (black colored) ranked based on fold change; red colored are memory related genes. **K**. Pathway analysis of common upregulated genes from j. **L**. 40x IF images of presynaptic marker, synapsin (magenta) and DAPI (blue). Scale bar 50 µm. **M**. Quantified mean intensity (n=9 / treatment, mean ± s.e.m, *unpaired t-test*, P*<0.05). **N**. Heatmap showing hierarchical clustering of z-score normalized gene expression counts of the union of DEGs (*P-adj* < 0.05) across all conditions **O.** KEGG pathway analysis per cluster.

We examined the impact of ACSS2 upregulation on transcription within the primary hippocampal neurons, via RNA-sequencing (RNA-seq). We assayed two distinct time points, at 9 days and 21 days *in vitro* (DIV) to capture gene expression changes during different phases of neuronal maturation. Comparing the two-time points, we found 7,587 differentially expressed genes (DEGs) (*P-adj* 0.05) (**Fig. 1G**). There was upregulation of mitochondrial, lysosomal, and senescence-related genes in the 21 DIV neurons (**Fig.1, G and H**, and Fig. S1C). In contrast, there was downregulation of synaptic genes, enriched in pathways related to axon guidance, glutamatergic and cholinergic synapses (**Fig.1, G and I**, and Fig. S1D)—these are critical neurotransmitter systems in hippocampal memory function and undergo degeneration and dysfunction in aging and disease (*27–31*). The results indicate a dynamic shift in the molecular landscape of neurons over time, predominantly affecting gene expression associated with synaptic integrity and function.

To gain insight into the impact of ACSS2, we assessed DEGs between AAV.ACSS2 and vector control at 9 DIV (4 days post-transduction) vs 21 DIV (16 days post-transduction). We identified 1,076 and 627 DEGs (*P-adj*<0.05), respectively (**Fig. S1, E and F**), and found upregulation of neuronal and synaptic pathways at both time points **(Fig. S1G**), whereas downregulation in RNA metabolism and stress response at 9 DIV and lipid metabolism at 21 DIV (**Fig. S1H**). DEGs that overlap these two time points revealed upregulation by AAV.ACSS2 of consensus signature genes associated with synaptic transmission and AMPA receptor activity (Mef2c, Syt1, Nlgn2, and others), learning and memory (Grin1, Shank1, Neto1, Shisa7, and others), (hypergeometric test p < 1.669e-21) (**Fig. 1, J and K**). One key upregulated gene in the consensus signature was Myocyte enhancer factor 2c (Mef2c) (**Fig. 1K**, left, bottom, green bar), implicated in cognitive resilience (*32, 33*); Mef2c upregulation confers protection against neurodegeneration (*32*). In contrast, we detected a smaller number of DEGs downregulated in common between the two time points (**Fig. S1I**). The number of commonly downregulated DEGs was not sufficient for Gene Ontology analysis (GO). However, among these genes was HtrA1 (high-temperature requirement serine peptidase A1), with important proteolytic degradation of pathological deposits (*34*) and whose inactivity is associated with tau and amyloid aggregation (*35–37*)—two major hallmarks of AD (**Fig. S1I**). Genes associated with AAV.ACSS2 were further examined in the context of synaptic location using the SynGO database (*38*), revealing genes associated with both the pre- and postsynaptic compartments (**Fig. S1J**). In particular, in hippocampal neurons we confirmed AAV.ACSS2-mediated upregulation of synapsin, a pre-synaptic marker, using immunofluorescence imaging (**Fig. 1, L and M**); synapsin controls release of neurotransmitters, including critical glutamate (*39*), with a crucial role in synaptic plasticity, which is a process fundamental to learning and memory (*40, 41*).

Distinct neuronal patterns of gene regulation over time and with AAV.ACSS2 were revealed by hierarchical clustering of z-score normalized DEGs (**Fig.1, N and O**). Neurons exhibited intrinsic reduction from 9 to 21 DIV in gene expression of nervous system development and axon guidance, which was countered by AAV.ACSS2 (clusters 2, 3 and 4) (**Fig.1, N and O**). Upregulation of mitochondrial genes (cluster 5), and proteasomal genes (cluster 6) became evident over time, which was also countered by AAV.ACSS2 (clusters 5 and 6) (**Fig.1, N and O**). This ACSS2-mediated reduction in mitochondrial and proteasomal genes may be attributed to a cellular state in which metabolic modifications are less necessary for maintenance of neuronal health and function, because neurons with AAV.ACSS2 are operating with improved homeostasis and overall health.

Further, we identified an intrinsic decrease at 21 DIV in expression of genes encoding histone modifying enzymes, including histone acetyltransferases 2A, 6A, and 6B (Kat2a, Kat6a, Kat6b), and histone deacetylases (HDAC 2, 4, and 6); again, these trends were countered with upregulated ACSS2 (**Fig. S1K**). Reduction in epigenetic and histone-modifying enzymes is a feature of aging and disease (*23–25, 42, 43*). Collectively, these data indicate that ACSS2 upregulation enhances and maintains expression of neuronal and synaptic genes over time in hippocampal neurons *in vitro*.

### AD-tau pathology diminishes synaptic gene expression and reduces H3K27ac in primary hippocampal neurons

Tau plays a prominent role in ADRD pathology with spreading of pathological tau across the brain, converting normal tau protein into the pathological hyperphosphorylated form (*20–22, 44, 45*). Given above insight gained from our transcriptomic analysis highlighting the potential of upregulated ACSS2 to improve synaptic gene expression, we investigated ACSS2 upregulation in promoting resilience against tau-associated pathology. We adapted an *in vitro* model of primary hippocampal neurons treated with human AD-tau (hAD-tau) (*46*) which models sporadic tau pathogenesis without tau overexpression (*20, 47, 48*).

First, we characterized gene expression changes with hAD-tau treatment alone, performing RNA-seq of hippocampal neurons treated with hAD-tau for 14 days. Comparing transcriptomic changes between hAD-tau and PBS treated (control) neurons identified 299 differentially expressed genes (*P-adj* <0.05) (**Fig. 2A**). Among significant hAD-tau-upregulated genes were apolipoprotein E (*APOE*), Adrenomedullin (*Adm*), and serpin peptidase inhibitor (*Serpina3n*); importantly, these genes are involved in AD pathogenesis (*49–51*). Further, the activator protein 1 (AP-1) superfamily members (JUN and FOS) were upregulated (**Fig. 2, A and B**), which regulate neuronal plasticity and activity, and are associated with aging (*52*). GO analysis of significantly hAD-tau-downregulated genes revealed pathways of synapse biology and synaptic membrane potential (**Fig. 2C**). Moreover, Mef2c was significantly hAD-tau-downregulated (**Fig. 2A**), also observed above as decreasing in neurons at 21DIV (**Fig. 1, J and K**). We queried the AD human brain gene expression data sets from Mayo Clinic (*53*) and from the Religious Orders Study and Rush Memory and Aging Project (ROSMAP) (*54*), which revealed dysregulation of our 299 hAD-tau DEGs (P<0.0001) in human AD patient brains (**Fig. 2D**). These data underscore that hAD-tau burden exerts a broad effect on the neuronal transcriptome that is captured in this *in vitro* tau spreading model system.

**Fig 2.**
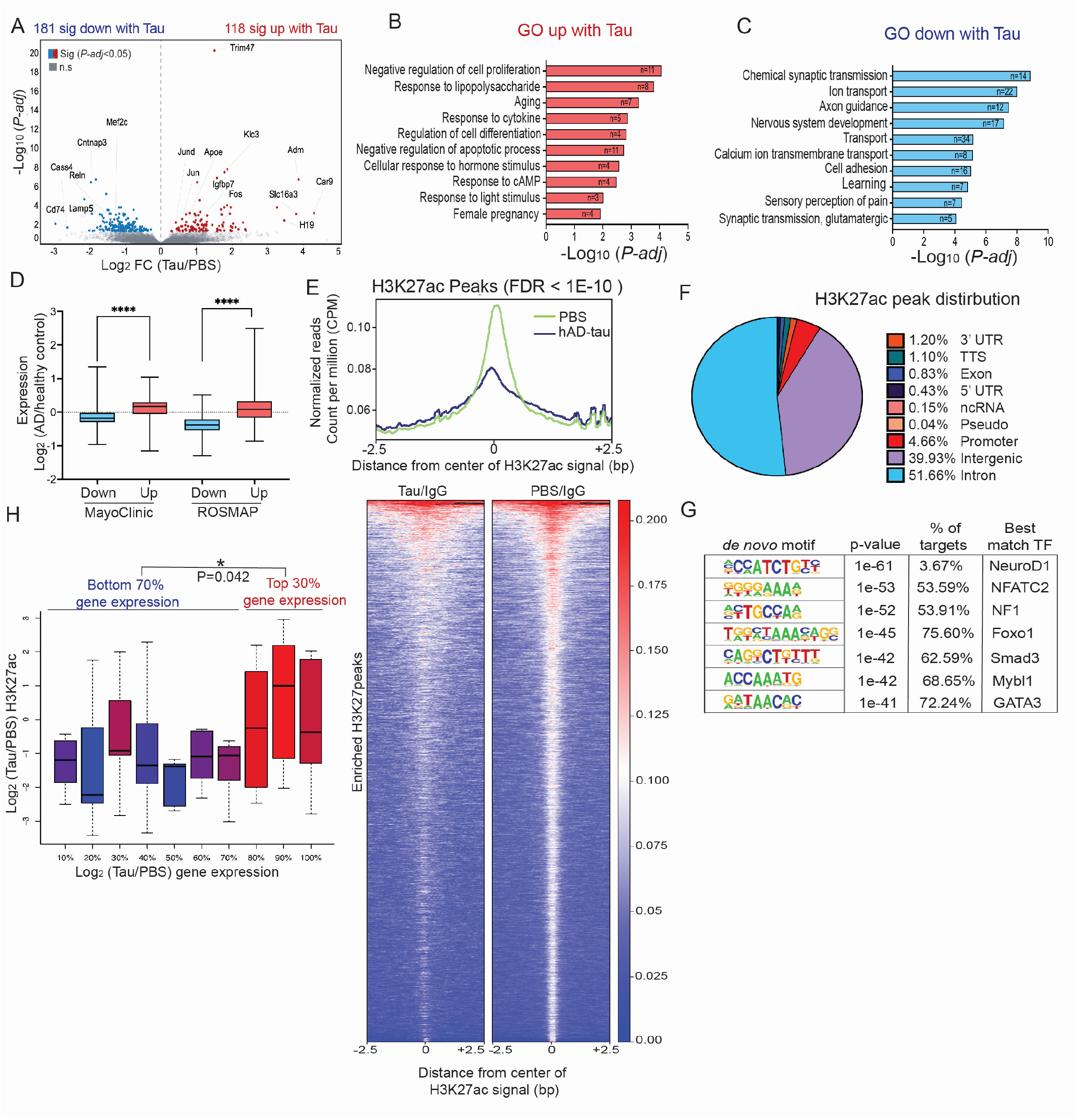
AD-tau induces global transcriptomic and epigenetic changes in neurons. **A**. Volcano plot of RNA-seq data from 21 DIV hippocampal neurons showing DEGs (*P-adj* 0.05) that are upregulated (red) or downregulated (blue) with hAD-tau treatment compared to PBS (control). All other genes are colored gray. Genes of interest are labeled. **B.** Top GO terms identified for upregulated, and **C.** downregulated significant DEGs of tau-stimulated neurons. **D**. Box plot represents log2 FC comparing AD patients to control for the 299 significant DEGs in neurons from Fig 2a. **E.** Profile of H3K27ac Cut&Run enrichment near protein coding genes. H3K27ac signals averaged from n=2 biological replicates per condition. Upper panels show the average profile around detected peak centers. Lower panels show read density heatmaps around the detected peak centers (9,649 differentially enriched H3K27ac peaks with FDR cutoff 1E-10). **F.** Distribution of differential H3K27ac peaks within the genome **G.** *De novo* Motif analysis showing enrichment of neuronal transcription factors at H3K27ac peaks. **H.** Genes with reduced expression show loss of H3K27ac levels. FDR cutoff 1E-10 of the H3K27ac differential peaks, Mann-Whitney non-parametrical test (P = 0.042) used for the comparison of H3K27ac levels between the bottom 70% of gene expression values and the top 30% gene expression values.

Transcriptome-correlated changes in H3K27ac levels were assessed via Cut&Run, which showed that hAD-tau treated neurons had significant reduction in global H3K27ac levels (**Fig. 2E**, upper). Using an FDR cutoff of 10E-10, we detected 9,649 differentially enriched peaks by contrasting hAD tau-treated with PBS-control cells and using matching IgG samples as background control (**Fig. 2E**, lower). Genomic distribution of these differential peaks preferentially appeared in the intronic (51.66%) and intergenic (39.93%) regions (**Fig. 2F**). DNA motif enrichment analysis of differential H3K27ac peaks identified binding sites of transcription factors NeuroD1 associated with neurogenesis (*55*), NFATC2 with a role in efficient axonal growth (*56, 57*), and NF1 associated with learning and cognitive function (*58*) (**Fig. 2G**). Further, downregulated genes correlated with loss of H3K27ac, while upregulated genes showed increased H3K27ac (**Fig. 2H**). Overall, H3K27ac levels and transcriptomic data largely showed the same direction of effect (**Fig. 2H and fig. S2A**). Further, hippocampal neurons showed significant reduction in ACSS2 protein levels with hAD-tau treatment, and modest reduction in H3K9ac levels (**Fig. S2, B to D**), indicating that changes induced by tau negatively impact ACSS2 expression. Together, these data reveal decreased expression of learning-related genes and upregulation of age-associated genes associated with reduced ACSS2 and histone acetylation (H3K27ac) in primary hippocampal neurons upon hAD-tau treatment.

### ACSS2 upregulation counters hAD-Tau induced transcriptome changes and enhances primary hippocampal neuronal resilience to pathology

We investigated whether ACSS2 upregulation can counter transcriptome changes induced by hAD-tau revealed in the previous section, via priming the isolated hippocampal neurons with AAV.ACSS2 prior to hAD-tau treatment (**Fig. 3A**). We assessed hAD-tau-induced transcriptome changes with and without AAV.ACSS2 at 9 and 21 DIV, corresponding to early and late stages of hAD-tau pathology (*59*) (**Fig. 3, A to D**), and carried out unsupervised clustering of DEGs compared to AAV.ACSS2 alone (**Fig. 3B**). Consistent with absence of aggregation at 9 DIV (*59*) there was a similar pattern of transcriptional changes induced by hAD-tau in hippocampal neurons with AAV.ACSS2 compared to control vector (**Fig. 3B**). However with increased tau pathology at 21 DIV (*46*), hippocampal neurons primed with AAV.ACSS2 altered the gene expression changes induced by hAD-tau (**Fig. 3B**). Specifically, AAV.ACSS2 globally altered the transcriptional response to hAD-tau, with 8 distinct gene clusters identified by hierarchical clustering, using standardized log fold-changes of hAD-tau with AAV.ACSS2 or vector compared to vector treated PBS controls (**Fig. 3B**). In clusters 4, 5, and 7, AAV.ACSS2 augmented the effect of hAD-tau compared to hAD-tau alone (**Fig. 3, B and C** and Fig. S3, I,J, and L). GO analysis of genes within clusters 4 and 5 revealed an enrichment of biological processes associated with neuron differentiation, synaptic function, and sterol biosynthesis, which were further enhanced with AAV.ACSS2 (**Fig. 3, C, D** and Fig. S3, I and J). Cluster 7 genes showed reduction of ER calcium ion homeostasis and apoptosis, and their expression was further reduced with AAV.ACSS2 (**Fig. 3, C and D**, and fig S3L). We infer that these clusters are a compensatory response to hAD-tau, since their expression was further facilitated in the same direction with AAV.ACSS2.

**Fig 3.**
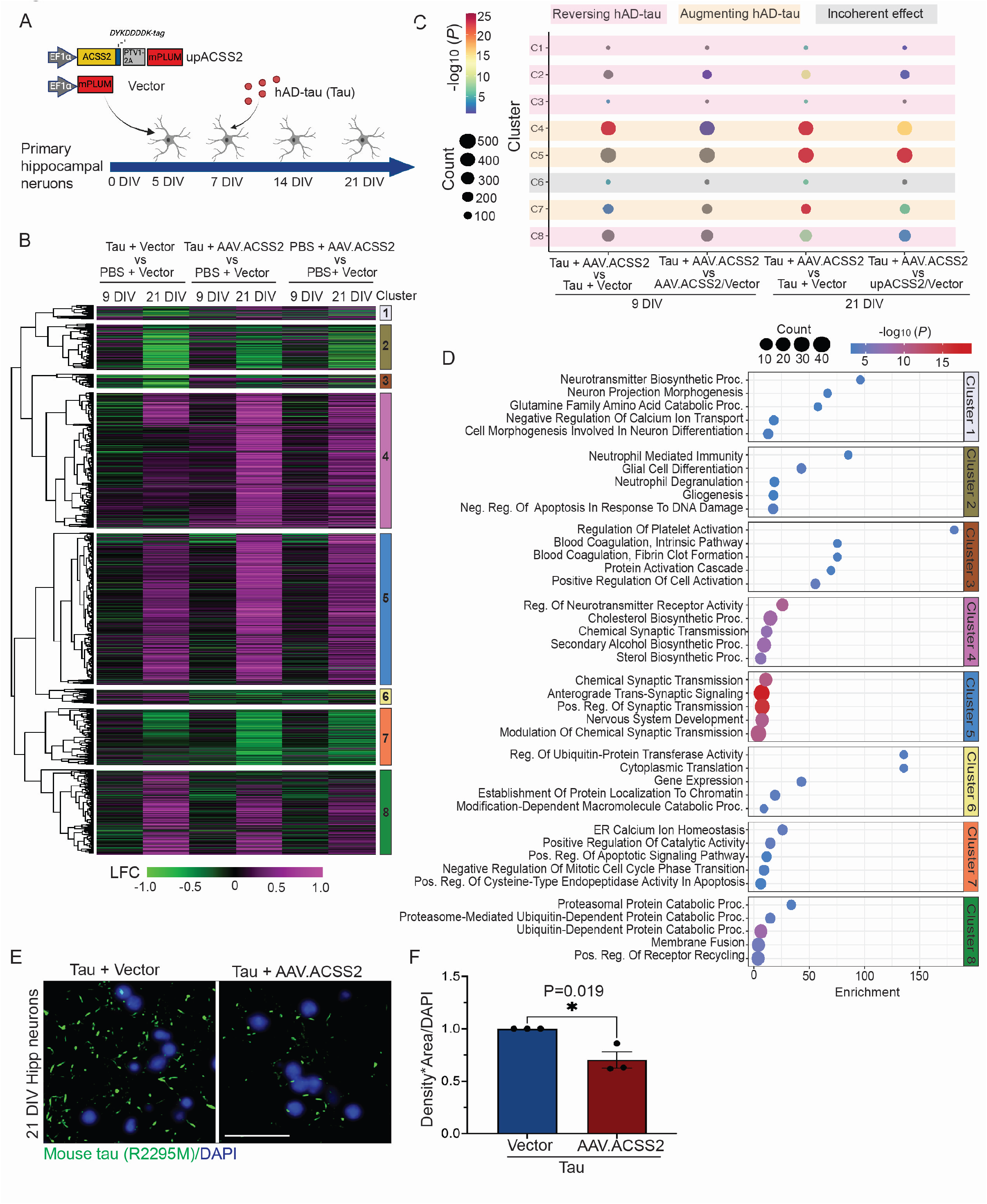
ACSS2 upregulation alters tau induced transcriptomic changes and enhances neuronal resilience to tau pathology. **A.** Schematic illustrating ACSS2 upregulation in hippocampal neurons treated with hAD-tau. **B.** Heatmap of 1800 DEGs (*P*-*adj* <0.05), using standardized log fold changes (LFC) of hAD-tau or AAV.ACSS2 (or both) against a background of PBS + vector controls **C.** Permutation test showing the summary of contrasts within clusters: bubble plot depicting -log10(p) values versus gene count (#genes). All analyses were conducted using permutation tests with the ‘coin’ package. **D**. KEGG pathway analysis of hierarchical clustering of DEGs. The size of dots for each pathway represents counts of enriched genes, and dot colors represent *P-adj* value. **E**. High content IF imaging of 21 DIV hippocampal neurons stained with mouse tau-specific anti-tau antibody (R2295M, green) and DAPI (blue), Scale bar 50 µm. **F.** Quantification: density x area normalized to DAPI. n=3 biologically independent samples. Average signal intensity was automatically quantified from 12 fields of view per well, 4-10 wells were used per biological replicate, mean ± s.e.m, *Unpaired t-test*, *P <0.05, and analysis was performed in a blinded manner.

In contrast, clusters 1,2,3, and 8 showed AAV.ACSS2 potential reversal of the hAD-tau effect: hAD-tau + AAV.ACSS2 DEGs were less pronounced than tau alone (hAD-tau + vector) and, interestingly, no significant difference compared to AAV.ACSS2 alone (PBS + AAV.ACSS2/ PBS + vector) (**Fig. 3, B and C**, and Fig. S3, F to H, and M). GO analysis of cluster 1 revealed increase in neurotransmitter synthesis and neurogenesis related pathways, which were downregulated with hAD-tau. Cluster 2 showed a reduction of genes related to glial cell differentiation and DNA damage response. Cluster 3 was associated with the regulation of platelet activation (**Fig. 3, C and D** and Fig. S3, F and H), perhaps as a mechanism to regulate inflammatory response (*60*) or to promote new synapses (*61, 62*). Cluster 8 was related to proteasomal functions and these genes were upregulated with hAD-tau, suggesting a compensatory mechanism aimed at clearing protein aggregates (*63–65*); at 21 DIV, expression of these genes was relatively diminished with AAV.ACSS2, suggesting a potential regulation of proteasomal pathways by AAV.ACSS2 (**Fig. 3, C and D** and fig S3M). ACSS2 upregulation did not show significant impact on genes in cluster 6 (**Fig. S3K**). We summarized the expression of these genes within each cluster using permutation test and FDR controlled Benjamini-Hochberg to validate significance and assess overall transcriptome effects of AAV.ACSS2 on hAD-tau (**Fig. 3C** and Fig. S3, F to M).

Based on these observations, we examined whether AAV.ACSS2 conferred resilience against tau pathology. We treated primary neurons with hAD-tau with/without AAV.ACSS2, and stained tau aggregates with anti-mouse tau antibody (R2295M; see Methods) after removal of soluble protein in the cell. Here, we observed significant reduction (P=0.019) of endogenous tau pathology in primary hippocampal neurons with AAV.ACSS2 (**Fig. 3, E and F**). Collectively these data suggest that ACSS2 upregulation counters AD-tau-induced transcriptome changes and contributes to resilience against tau pathology.

### Pre-symptomatic increase of ACSS2 levels in mouse AD genetic model rescues Tau-induced memory decline, plasticity, and pathology

Tau pathology is a major driver of cognitive dysfunction in ADRD (*21*), exerting its most detrimental effects within the hippocampus, which is crucial for memory. As tau pathology advances, synaptic function required for cognition and memory is disrupted (*20, 45, 66*). Our *in vitro* data indicated that upregulated ACSS2 countered effects of Tau spreading, mitigating transcriptional changes and tau pathology itself. To extend these data *in vivo*, we determined whether AAV.ACSS2 could increase neuronal resilience to tau-induced memory decline of the P301S (PS19) mouse, a transgenic mouse model of tauopathy with age-associated spatial memory deficits (*67, 68*). Animals were treated via bilateral stereotaxic injection into the dHPC (dorsal hippocampus) with the PHP.eB AAV.ACSS2 or vector control (**Fig. 4A**). The PHP.eB serotype AAV predominantly transduces NeuN+ neurons and thus minimizes potential heterogeneous effects of AAV injection (*69–71*). Mice were injected at 2.5 months to assess whether early intervention can enhance resilience to disease-associated perturbation, which emerges at ∼8-9 months (*72, 73*).

**Fig 4.**
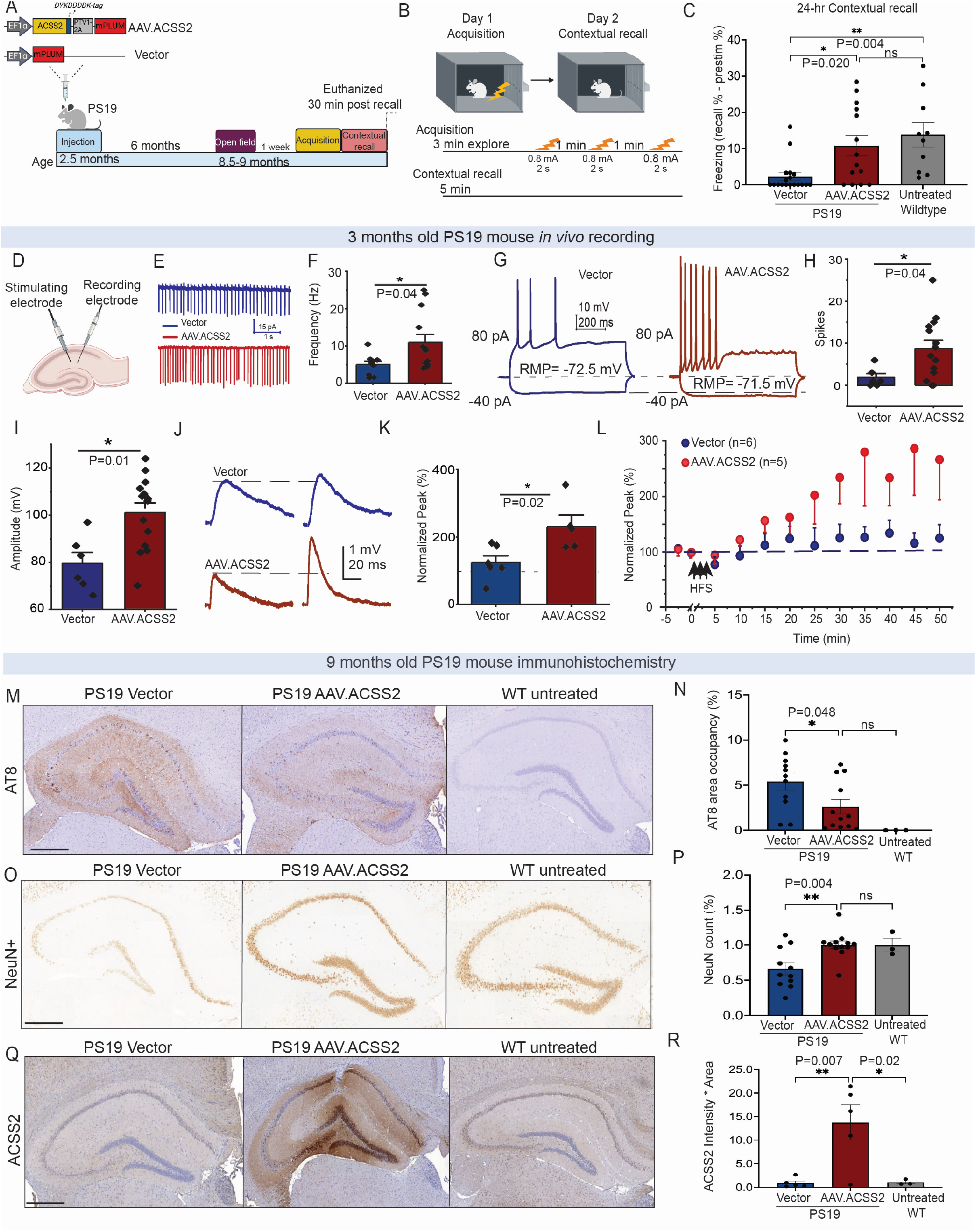
Pre-symptomatic enhancement of ACSS2 maintains resilience to tau-induced memory decline, synaptic plasticity, and transcriptomic changes. **A**. Schematic of the experimental design **B**. and fear conditioning. **C.** Contextual memory recall (One way-ANOVA, *P<0.05, **P<0.01, mean ± s.e.m.; n=18 PS19 vector, n=14 PS19 AAV.ACSS2, n=10 untreated wildtype). The experiment was conducted in three independent batches. **D**. Electrophysiology recording. **E**. Spontaneous action potential firing from CA1 pyramidal neurons. **F**. Summary of firing frequency. **G**. Typical single traces of the voltage membrane response to hyperpolarizing current and evoked action potential firings during the application of depolarizing current with RMP. **H**. Summary of the number of spikes evoked by 80 pA of injected current. **I**. Summary of the amplitude of the evoked action potentials. **J.** Typical traces of EPSP recorded in CA1 before (left) and 45 min after (right) LTP induction. **K.** Summary of the normalized EPSP amplitude 5 min before LTP induction and 45 min after LTP induction. **L.** Summary of the normalized EPSP amplitude vs. time. Three HFS (1s of 100 Hz repeated 3 times with intervals of 20s) induced LTP. Gray dashed lines indicate baseline levels of the EPSPs before HFS. **M.** IHC analysis of tau pathology (AT8), **N**. quantified, **O**. Neuronal survival (NeuN), (vector n=12, AAV.ACSS2 n=12 wildtype untreated n=3), **P**. quantified, and **Q.** ACSS2 (vector n=5 AAV.ACSS2 n=5 wildtype untreated n=3), **R**. quantified. *One-way ANOVA*, mean ± s.e.m, *P<0.05, **P<0.01. Each dot represents one mouse. All analyses were performed in a blinded manner.

In blinded behavioral studies, anxiety levels and spatial memory were examined at 9 months of age using the open field test and contextual fear conditioning (FC). There were no differences in anxiety-related behavior between the AAV.ACSS2 and control PS19 mice (**Fig. S4, A to D**) and, importantly, at 9 months the PS19 mice displayed full mobility with no motor deficits during open field test (**Fig. S4, E to H**). Moreover, there were no differences in acquisition of fear memory between the AAV.ACSS2 and control PS19 mice (**Fig. S4I**). In contrast, the AAV.ACSS2 PS19 mice exhibited significant increased fear memory manifested by freezing behavior in 24-hour contextual memory recall compared to the control group (**Fig. 4, B and C**). These findings indicate that ACSS2 upregulation enhances consolidation and long-term retention of contextual fear memory of PS19 mice.

Given the deficits in synaptic dysfunction and long term potentiation (LTP) in the PS19 mouse (*68, 72*), we hypothesized that memory improvement of ACSS2 upregulation may result from enhanced LTP. LTP is a primary synaptic plasticity change required for memory storage and retrieval, and its dysfunction generally underlies memory loss (*74–78*). Neuronal firing patterns induce changes in synaptic plasticity that can selectively strengthen or weaken neuronal networks (*79, 80*). In neurodegenerative diseases involving tau pathology, a subpopulation of neurons is reduced in intrinsic excitability (*81*). To examine LTP, we injected PS19 mice with AAV.ACSS2 or empty viral vector as before (**Fig. 4A**) and assessed background excitability at 1-month post injection (mpi) of CA1 pyramidal neurons (**Fig. 4D**), which are central to consolidation and retrieval of hippocampal-dependent memories (*75*). Using freshly cut brain slices, spontaneous action potential firing was recorded for 5-10 minutes. We found that spontaneous firing in CA1 pyramidal neurons from AAV.ACSS2 mice (11.0 ± 7.7 Hz, n=13) was statistically faster (**Fig. 4E**, P= 0.04) than firing from vector-control mice (5.0 ± 2.9 Hz, n=9) (**Fig. 4, E and F**). We then performed whole-cell recording of neurons in brain slices to further examine electrophysiological properties of the neurons. To assess differences in active and passive electrophysiological properties of CA1 neurons between AAV.ACSS2 and control, we examined neuronal response to a hyperpolarizing current (injected −40 pA) and to a depolarizing current (injected 80 pA) under current clamp mode (**Fig. 4, G and H**). We found that CA1 pyramidal neurons from AAV.ACSS2 mice fired more action potentials (8.8 ± 1.9 Hz, n=14) than control mice (1.8 ± 1.0 Hz, n= 6) (**Fig. 4G**, P=0.04). Further, the amplitudes of the evoked action potentials were also significantly greater with ACSS2 upregulation (101.2 ± 4.1 mV, n= 14) than control (79.6 ± 4.7 mV, n= 6) (**Fig. 4I**, P=0.01). These findings indicate that the excitability of CA1 pyramidal neurons was significantly elevated with ACSS2 upregulation.

Next, we examined LTP induction (**Fig. S4J**), first determining the input-output relationship of the basal synaptic strength in each CA1 pyramidal neuron to evaluate the range of whole-cell excitatory post-synaptic potentials (EPSPs) achievable in response to stimulation. We found no significant differences in EPSPs between control and AAV.ACSS2 mice (**Fig. S4, K and L**). Following LTP induction by high-frequency stimulation (HFS), postsynaptic currents were significantly greater in the AAV.ACSS2 mice (P=0.01, 231.2 ± 33.4% vs baseline, n=5, paired t-test) compared to control mice (P=0.3, 125.0 ± 20.2% vs baseline, n=6, paired t-test) (**Fig. 4, J and L**). This difference in LTP formation indicates improved CA1 pyramidal neuron function with AAV.ACSS2 mice.

Neuropathological protein accumulation in AD disrupts the balance of inhibitory and excitatory synaptic transmission, propagating neuronal dysfunction (*20, 81*). To determine whether improved electrophysiological features and memory of PS19 mice with AAV.ACCS2 correlated with a change in tau pathology, we immunostained for pathological tau (AT8 antibody) in the hippocampus of 9 month PS19 mice, revealing notable reduction in tau pathology with AAV.ACSS2 (P=0.048) (**Fig. 4, M and N**; control in **Fig. 4, Q and R**) and associated with significant improvement in NeuN+ immunostaining (P=0.004) (**Fig. 4, O and P**) indicating reduced neurodegeneration. These observations extend our *in vitro* findings in isolated neurons (see **Fig. 3, E and F**), here showing that AAV.ACSS2 protects against pathological tau accumulation. Overall, the electrophysiology and immunohistochemistry reveal a strongly protective role exerted by ACSS2 upregulation against tau-induced detrimental effects *in vivo*.

### Global changes of CpG DNA methylation in AD mice are reversed with ACSS2 upregulation

We assessed molecular features of AAV.ACSS2 in the PS19 mice. First, we analyzed gene expression associated with improved memory via RNA-seq of the dorsal hippocampus (dHPC) of mice euthanized 30 min post contextual recall (**Fig 5A**) given strong dependency of contextual recall on the dHPC region (*82, 83*). RNA-seq was performed on 1 and 6 month post injection (mpi) of AAV.ACSS2 or control vector, and mice of both time points displayed improved memory (**Fig. S5A**). We found increased expression of synaptic genes with AAV.ACSS2 at 1 mpi (**Fig. 5B** and **Fig. 5D**), pointing to an early and robust effect of ACSS2 on synaptic function and indicating molecular mechanisms underlying the improved LTP and memory observed in PS19 mice (see **Fig. 4C** and **Fig. 4L**). Analyses at 6 mpi revealed changes in fundamental cellular processes, including metabolic and ribosomal genes (**Fig. 5C and D**). These data thus highlight a dynamic temporal profile of gene expression changes, with early synaptic changes followed by alteration in broader cellular processes that likely support long-term memory enhancement. The impact of AAV.ACSS2 on synapses was most pronounced relatively early after injection (by 1 month) leading to strengthening of synapses and LTP and, even after synaptic gene expression returns to baseline levels, memory-enhancing effects persisting over time (until 6 months).

**Fig 5.**
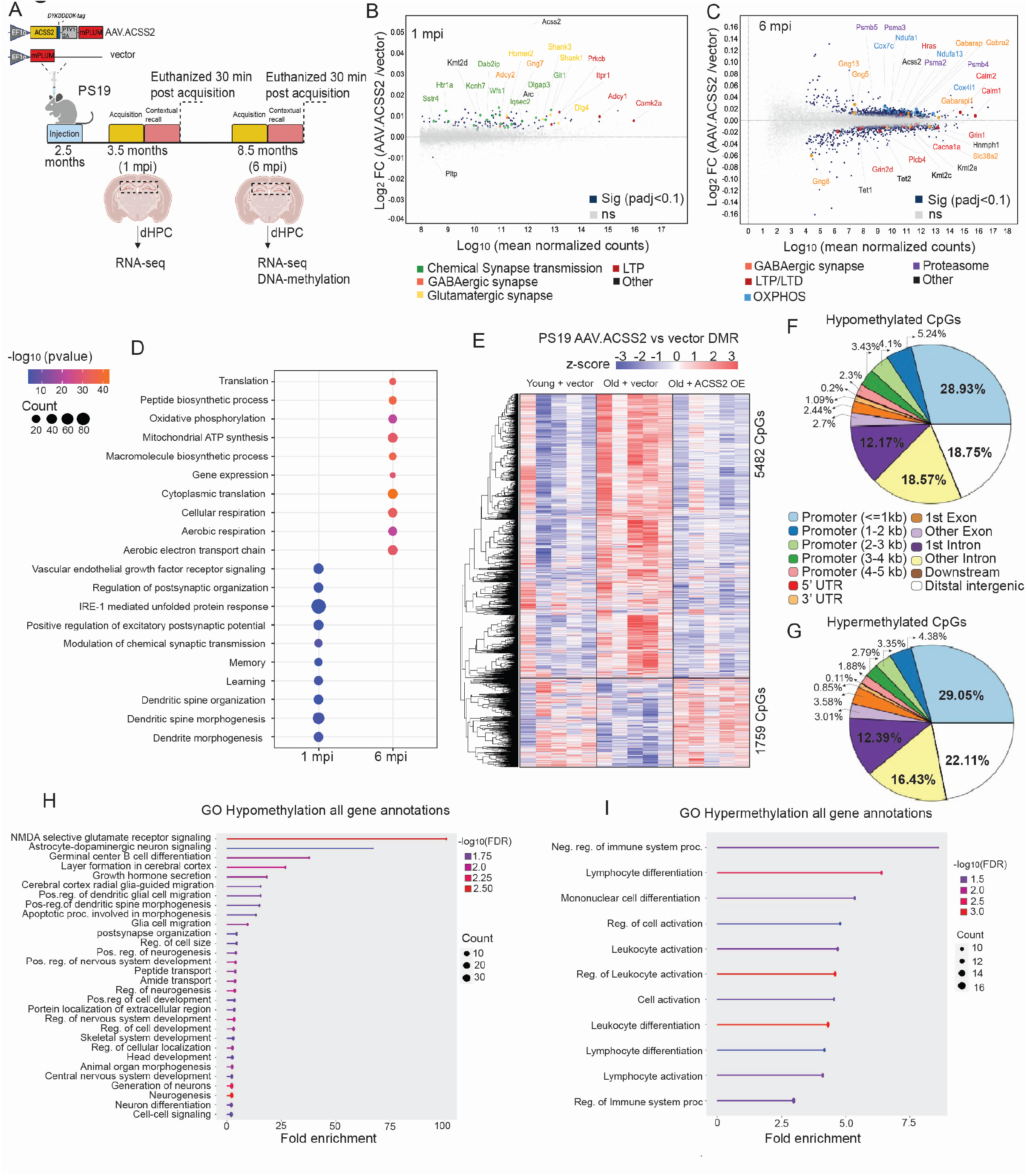
Global changes to CpG DNA methylation in AD mice are reversed with AAV.ACSS2. **A.** Schematic illustrating the experimental design in PS19 mice. **B.** MA plot of RNA-seq data from bulk dHPC tissue from 3 months-old (1mpi), and **C.** 8.5-9 months old (6 mpi) PS19 mice. Significant DEGs (*P-adj* < 0.1) are highlighted in dark blue, while non-significant genes are displayed in gray. Significant genes of interest are color-coded according to their respective KEGG pathways. n=5 per time point AAV.ACSS2, n=5 per time point vector control). Data was analyzed using (RUVr) and regressing to the contextual recall score. **D.** GO analysis shows enrichment of synaptic and memory related genes and metabolic genes 1 mpi and 6 mpi, respectively. **E.** Heatmap representing z-score scaled single CpGs using whole hippocampi. **F.** Distribution of hypomethylated (5482 CpGs), and **G.** hypermethylated (1759 CpGs). **H.** GO enrichment analysis of hypomethylated, **i.**, and hypermethylated CpGs in gene promoters.

Next, we assessed potential persistent epigenetic effects, given the remarkable long-term impacts following a single dose of AAV.ACSS2, to reverse disease and rescue memory deficits 6 months post-injection in the PS19 mice. While histone acetylation is integral to chromatin alterations regulated in part by ACSS2, the sustained synaptic and memory enhancement suggested involvement of additional layers of long term epigenetic programming, such as DNA methylation (*84*). Moreover, there is strong association between increased DNA methylation, elevation of pathology, and decline in global cognitive function in AD patients (*85–88*), indicating a role of these epigenetic mechanisms in driving AD. Therefore, we investigated whether ACSS2 upregulation may counteract tau influence on DNA methylation. We found that CpG methylation of 9-month-old PS19 mice with AAV.ACSS2 closely resembled young (3-month) PS19 mice (**Fig. 5E** and **Fig. S5B**), compared to 9-month-old PS19 control mice. These findings suggest reversal by AAV.ACSS2 of CpG DNA methylation changes induced by aging of PS19 mice. Further, hypo- and hypermethylated CpG regions were enriched in promoter and intronic regions relative to the distribution of CpGs across these features on the whole array (**Fig. 5, F and G**). Next, we assessed the correlation between DNA methylation (associated with gene silencing) and RNAseq. We found a weak correlation between increased DNA methylation and gene silencing, assessing all DNA methylated regions (DMRs) or DMRs overlapping with promoters (**Fig. S5, C and D)**. Considering the stability of DNA methylation and the dynamic nature of gene expression, our analysis suggests that additional epigenetic chromatin regulatory mechanisms are involved in maintaining the long term gene expression patterns. We also point out that the CpG array is very limited in assessing the entirety of DNA methylation. Despite these limitations, GO analysis of the AAV.ACSS2-associated hypomethylated CpGs (5482 single CpGs) in promoters revealed genes associated with NMDA glutamate receptor signaling, astrocyte-dopaminergic neuron signaling, and neurogenesis (**Fig. 5H**). GO analysis of the AAV.ACSS2-associated hypermethylated promoter regions (1759 single CpGs) showed enrichment of immune system-related pathways (**Fig. 5I**). Together, these data indicate an indirect role of ACSS2 in regulating DNA methylation leading to long term epigenetic memory, and a dynamic interplay between ACSS2 and the broader cellular machinery governing gene expression relevant to improved electrophysiology and memory.

### ACSS2 upregulation prevents age-associated cognitive decline

Age is the dominant risk factor for ADRD. The remarkable outcomes of AAV.ACSS2 in preventing tau pathology and associated memory decline, coupled with reversing the DNA methylation patterns hold profound implications for potential rejuvenation of the aging brain and mitigating this primary risk factor. Hence, we investigated whether AAV.ACSS2 could protect against normal age-associated memory decline, in the absence of underlying pathology or disease.

To assess the impact of ACSS2, we used retro-orbital injection of AAV.ACSS2 in wild type adult (10-11 months) to assess both molecular and memory impacts. Retro-orbital injection of PHP.eB produces effective transduction of the CNS with minimal off-target transduction of the liver (*69, 71*). To capture early gene expression changes, we euthanized mice two weeks following ACSS2 vector injection, harvesting the dHPC 30 min after the acquisition of FC (**Fig. 6A**). RNA-seq analysis of AAV.ACSS2 compared to vector control revealed 328 upregulated and 40 downregulated genes (*P-adj* <0.1) and 199 upregulated and 11 downregulated genes (*P-adj* <0.05) (**Fig. 6B**). Genes increased in expression represented consensus signature genes associated with neuron projection and axons, and canonical genes involved in longevity-regulating pathways (e.g. Foxo3, Irs2, Mtor), circadian clock (e.g. Per1, Kcnj6, Cacna1h), and chromatin-modifying enzymes (e.g. EP300, EP400, Kat6a, Kmt2a, Jade2) (**Fig. 6C**).

**Fig 6.**
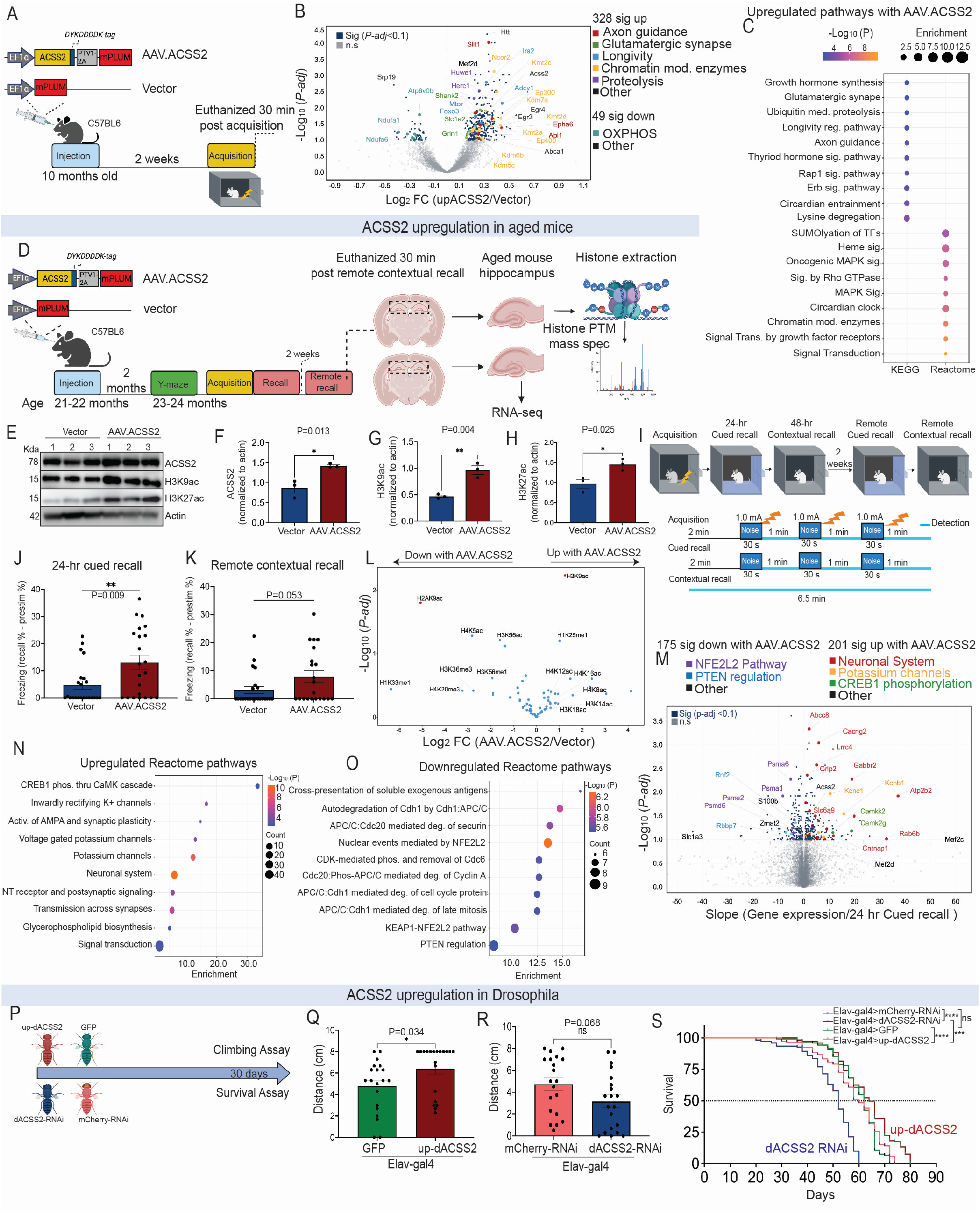
ACSS2 upregulation prevents age-associated cognitive decline. **A.** Schematic of experimental design **B.** Volcano plot of RNA-seq data from dHPC. Significant DEGs (*P-adj* < 0.1) are highlighted in dark blue, while non-significant genes are displayed in gray. Significant genes of interest are color-coded according to their respective pathways (AAV.ACSS2 n=6, vector control n=5). **C.** Pathway analysis. **D.** Experimental design in aged mice. **E.** WB of whole mouse hippocampus. **F**. Ratio of ACSS2, **G.** H3K9ac, **H.**, and H3K27ac to actin (n=3; unpaired *t-test*, mean ± s.e.m, *P<0.05, **P<0.01). **I**. FC schematic **J.** 24-hr contextual recall reflected as fold-change over pretone interval. **K**. Remote contextual recall reflected as percent time freezing, averaged over the entire recall period (5 min). AAV.ACSS2 n=22, vector n=23, unpaired *t*-test, mean ± SEM, **P<0.01. Experiment was performed in 4 independent batches and analysis was performed in a blinded manner. **L.** Volcano plot showing histone PTM differences from whole hippocampi. Significant (P<0.05, unpaired *t-test*) histone acetylation changes are highlighted in red. **M.** Volcano plot of RNA-seq data from aged mice hippocampus (n=12 AAV.ACSS2 and n=12 vector, Data analyzed using RUVr and regressing to the 24-hour cued recall score), **N.** upregulated, and **O.** downregulated pathways **P**. Schematic of experimental design. **Q.** Climbing assay in drosophila with up-dACSS2, **R.** or ACSS2 RNAi, unpaired *t-test,* *P<0.05. s. Kaplan-Meier Survival analysis, n=487 per condition.

We then investigated impacts of ACSS2 upregulation on age-associated cognitive decline. Aged mice (21-22 months old) were injected via retro-orbital injection (**Fig. 6D**) with AAV.ACSS2 and control vector. ACSS2, H3K9ac, and H3K27ac levels were elevated in the hippocampus at two months post-injection (**Fig. 6, E to H**) consistent with sustained expression of ACSS2. In a blinded behavioral study, we examined short- and long-term memory (**Fig. 6I**). There was no effect of AAV.ACSS2 on short-term memory by Y-maze assay (**Fig. 6S, A and B**), and no difference in baseline freezing behavior during the acquisition phase in the FC assay (**Fig. S6C**). In contrast, there was notable increase in freezing behavior in AAV.ACSS2 mice with 24-hour cued recall (**Fig. 6J**), but not in 48-hour context recall (**Fig. S6D**) possibly due to extinction learning during the 24-hour cued recall (*89*). Consistently, there was an attenuation of freezing behavior with AAV.ACSS2 in remote cued recall (**Fig. S6E**), suggesting learned dissociation between fear response and stimulus (cue) (*90*). Interestingly, there was improvement in remote (2 weeks) context recall with AAV.ACSS2 (**Fig. 6K**), which could result from strengthening of excitatory connections between prefrontal cortex neurons (*91*); indeed AAV.ACSS2 was detected in the aged brain prefrontal cortex (**Fig. S6F and G**). Taken together, these data indicate that ACSS2 enhances age-associated long-term memory.

Following the behavioral assays in the same aged mice cohorts, we examined activity-dependent global histone modifications and gene expression in the hippocampus. Mice were euthanized 30 min after the remote contextual recall test to assess ACSS2-dependent activity-induced changes. Consistent with immunoblotting results and *in vitro* data (**Fig. 1, D and E**) profiling of histone post-translational modifications by mass spectrometry revealed significant upregulation in H3K9ac levels (**Fig. 6L**). Concurrently, we analyzed gene expression changes in aged mice with AAV.ACSS2, analyzing 12 mice per treatment group to provide a comprehensive transcriptional profile of aged mice. We note that the mice were randomly selected and included representatives from all four independent batches ensuring that we captured a diverse range of responses in the FC assay. We first confirmed that AAV.ACSS2 treatment led to increased ACSS2 transcript levels of each mouse used for the RNA-sequencing (**Fig. 6H**). Transcription analysis revealed increased expression of consensus signature genes associated with the neuronal system (e.g. Gabbr2, Abcc8, Cam2g, Comt, Cacng3), potassium channels (e.g. Kcnc1, Kcnj2, Kcnj10, Kcnj11), CREB1 phosphorylation (e.g. Camkk2 and Camk2g); in contrast there was downregulation of proteasomal genes (e.g. Psma1, Psma2, Psma6, Psma7, and Psmd6) (**Fig. 6, M to O**).

Given the memory enhancement in aged mice (**Fig. 6, J and K**) and increased expression with AAV.ACSS2 of longevity-related pathways in the dHPC of adult wildtype mice (**Fig. 6, B and C**), we reasoned that increased ACSS2 may impact health span and/or lifespan. Indeed, cognitive decline is typically associated with increased mortality rate (*92–96*). To investigate the physiological effect of altered neuronal ACSS2 levels, we utilized *Drosophila melanogaster*, as a powerful model system for aging (*97, 98*). We employed a neuronal driver, *elav*-GAL4, to generate a transgenic line that expressed *Drosophila* ACSS2 (AcCoAS) in all neurons (Elav-Gal4>dACSS2) with GFP as a control line (Elav-Gal4<GFP). In parallel we reduced ACSS2 levels within neurons (Elav-Gal4<dACSS2-RNAi), with mCherry siRNA knockdown serving as a control (Elav-Gal4< mCherry-RNAi) (**Fig. 6P**). Consistent with our hypothesis, upregulating neuronal ACSS2 significantly improved locomotor activity (**Fig. 6Q**) whereas ACSS2 knockdown reduced average locomotor activity (**Fig. 6R**). Notably, these effects manifested in aged flies (30 days) and not in younger flies (14 days) (**Fig. S6, I and J**). We determined whether improved health span is accompanied by increased longevity and lengthened lifespan, and, indeed, while knockdown of neuronal ACSS2 shortened lifespan, upregulation of ACSS2 extended lifespan (**Fig. 6S**). Taken together, these findings underscore a pivotal role of ACSS2 in neuronal function and potential beneficial impact on overall organismal health and lifespan.

## Discussion

Epigenetics, due to its dynamic yet enduring nature, provides a promising therapeutic avenue for mitigating age- and disease-related changes. The progression of aging in the brain is associated with altered epigenetic modifications within neurons, rendering them susceptible to disease (*5, 7, 11, 21, 26, 29*). It is widely acknowledged that molecular alterations associated with ADRD occur decades before onset of recognizable cognitive dysfunction and behavioral symptoms (*99*). The manifestation of symptoms becomes evident only as the aging process unfolds. Given the current lack of highly effective therapies for ADRD (*100*), most of which are primarily focused on reducing proteinopathy and are often associated with severe side effects, coupled with the modest impact on memory improvement (*101–103*), there is compelling need for new strategies. We reason that maximum benefit may be attained with a different approach that targets physiological dysfunction upstream of both protein pathology and the core impairment—cognitive dysfunction.

In this study, we carried out comprehensive investigation of ACSS2, a metabolic-epigenetic enzyme, as an approach to ameliorate age- and disease-associated cognitive decline. We demonstrate, via ACSS2 upregulation, enhancement of cognitive longevity and resilience against both aging and tauopathies. Our evaluation of molecular changes induced by ACSS2 upregulation in cultured hippocampal neurons revealed significant increase of gene expression pathways associated with synaptic function and neurogenesis processes that normally decline over time (**Fig 1G**). These findings underscore the potential of ACSS2 upregulation to preserve and enhance neuronal function. Conversely, we found that tau pathology suppressed expression of synaptic genes while amplifying aging signatures. Further revealing the beneficial effects of ACSS2, we showed that ACSS2 upregulation induces transcriptional changes that confer neuronal resilience to pathogenic accumulation of AD-tau (**Fig. 3, B and E**). Moreover, we observed that neurons with upregulated ACSS2 were protected from pathogenetic tau accumulation (**Fig. 3, D and E**).

Expanding on positive effects of ACSS2 upregulation, we conducted experiments in two distinct mouse models, including genetic and wildtype strains and varying age groups. Across these diverse conditions, we observed an overarching pattern of gene expression consistent with the *in vitro* model, showing, with upregulation of ACSS2, increase of synaptic genes, neurogenesis, and regulation of longevity-related pathways in aging and diseased cellular environment (**Fig. 5, B to D** and **Fig. 6, B and C**). This consistency indicates that the observed changes hold broad applicability in mitigating the effects of cognitive aging and neurodegenerative diseases. Our findings that boosting ACSS2 protects from tau-associated pathology and toxicity are highly impactful for an approach to mitigate disease-associated situations, given the prominence of tau in Alzheimer’s and other tauopathies.

Notably, in our study, a single injection of ACSS2 upregulation vector into PS19 (tauopathy) mice improved tau-induced cognitive dysfunction at 6 months post-injection (**Fig. 4, A to C**). These mice exhibited comparable baseline freezing behavior, and no confounding reduction of locomotor activity or increased anxiety, reinforcing that improvements in contextual fear conditioning were genuine effects of better memory, and, further, do not have undesirable side effects.

Following injection of AAV.ACSS2, we observed increased synaptic gene expression in the dHPC at the early timing of 1 month (**Fig. 5B**). In agreement with this gene expression profile, we detected significant enhancement in LTP at the 1-month injection mark (**Fig. 4K**), indicating strengthening of synapses (*104*). These synaptic enhancements appeared enduring as the animal aged, translating into improved memory observed in the fear conditioning behavioral studies (**Fig. 4C**). These data support the notion that early synaptic strengthening protects neurons from later degeneration (*28, 105*).

Consistent with the *in vitro* primary neuron culture model (**Fig. 3, E and F**), we also observed substantial reduction in tau pathology in these mice with ACSS2 upregulation (**Fig. 4, L and M**). In the *in vitro* model, ACSS2 upregulation protected healthy neurons from exogenous introduction of tau, via spreading and toxicity, which occurs in sporadic AD; in contrast, the *in vivo* model underscores a protective role of ACSS2 upregulation where the disease-causing genetic mutation is present throughout life. Given the role of tau in various neurodegenerative diseases such as AD, frontotemporal dementia (FTD) (*21*), and Parkinson’s disease with dementia (PDD) (*106*), reducing tauopathy through ACSS2 upregulation holds promise for a broad spectrum of therapeutic applications.

We investigated DNA methylation, which correlates with biological aging (*107, 108*). Examining the CpG DNA methylation patterns in adult (9 months) PS19 mice with upregulated ACSS2, revealed an intriguing association of ACSS2 with alterations of tau-induced DNA methylation (**Fig. 5E**). While both DNA methylation and histone acetylation are relevant to gene expression, the mechanism through which upregulated ACSS2 reduces tau-induced DNA methylation, and whether this process is directly linked to histone acetylation or operates independently, all require further exploration.

Considering the impact of ACSS2 upregulation in PS19 mice, we subsequently investigated an effect of ACSS2 in age-associated memory decline to focus on potential impact of ACSS2 in the context of aging alone, and to explore potential of ACSS2 as a rejuvenating intervention independent of confounding features associated with ADRD. Our findings revealed significant enhancement in cued and remote contextual recall (**Fig. 6, J and K**), suggesting that ACSS2 upregulation has capacity to improve long-term memory apart from pathology. We profiled gene expression changes correlating with improvement in cued memory recall in aged mice using a large cohort that represented varying degrees of memory improvement with ACSS2 upregulation. This approach enabled comprehensive characterization in the aging mouse of molecular changes associated with upregulated ACSS2 (**Fig. 6M**), and thus add to the robustness and reliability of our findings. Lastly, using *Drosophila* as a powerful model system for aging, we found elevated neuronal ACSS2 led to significant improvement in motor function and lengthening lifespan (**Fig. 6, Q to S**).

In conclusion, our study marks a significant milestone in understanding molecular features of ACSS2, but, importantly, also underscores broader implications of metabolic epigenetics in shaping cognitive outcomes. ACSS2 represents a novel, pleiotropic strategy to promote broad neuroprotection against the complex and diverse pathology observed in ADRD. Increased ACSS2 provides robust protection of neurons from toxic tau in these models, which clinically has been a difficult therapeutic target (*100, 109, 110*). The magnitude of the anticipated rise in individuals suffering from AD and age-related dementia, coupled with the predicted economic burden of over a trillion dollars in healthcare in the coming decades (*111*), underlines the urgent need to address these issues comprehensively. As traditional therapeutic approaches that target neurodegenerative protein aggregations fall short in efficacy, a strategy that instead incorporates epigenetic interventions holds promise in reversing age- and disease-related changes.

## MATERIALS AND METHODS

### Study Design

The aim of this study was to investigate whether enhancing ACSS2-dependent chromatin processes could protect neurons against gene dysregulation associated with disease and aging, thereby mitigating cognitive decline. Firstly, we evaluated ACSS2-dependent and AD-tau-dependent transcriptional changes in neuronal cultures. Secondly, we examined the effects of upregulating ACSS2 in ameliorating phenotypes in a tau-dependent neurodegenerative model. Thirdly, we explored the role of ACSS2 in rejuvenation in aged models. Male mice were utilized for all studies and were age-matched to ensure similarity between treatment and control groups. Specific sample sizes for experiments were detailed in the figure legends. Animal allocation to experimental groups was randomized, and experiments were conducted in a blinded manner. FC data was collected from aged mice across four independent batches and for PS19 mice across three independent batches. In FC assays with aged mice, two vector-treated aged mice were excluded based on Mahalanobis distance. No mice were excluded from the Open Field test and Y-maze. Furthermore, no PS19 mice were excluded from behavioral assays.

### Statistical Analysis

All experiments were conducted on a minimum of 3 biological replicates unless otherwise indicated. For primary neuron culture, biological replicates were defined as independent batches of cell culture. Statistical confidence was evaluated for Cut&Run peaks (FDR < 0.01) and upper quartile normalization was conducted for RNA-seq. All imaging analyses (IF, ICC, IHC) were conducted in a blinded manner. High-content imaging was performed on 3 independent experiments, and image quantification of the in vitro pathology was performed on random-field images. For IHC, the whole hippocampus was quantified. Significant differences were followed by Fisher’s least significant difference test when appropriate. Pairwise comparisons were made using Student’s t-test after testing for normality and equal variance. The threshold for statistical significance was considered α = 0.05 for all in vitro assays and drosophila models, and α = 0.1 for all mouse experiments. All nonsequencing statistical analyses used Prism GraphPad software to compute significance.

## Acknowledgments

We thank Andrew Davis (Sanford Burnham Prebys Medical Discovery Institute) for technical advice and assistance with the DNA-methylation array sequencing. The data generated at the UC San Diego IGM Genomics Center utilized an Illumina NovaSeq 6000 that was purchased with funding from a National Institutes of Health SIG grant #S10 OD026929. PHP.eB viral vectors and packaging were generated with the University of Pennsylvania Vector Core. We thank Lakshmi Changolkar (UPenn) for the material and logistical support. Behavior procedures were performed with The Neurobehavior Testing Core at the University of Pennsylvania/Institute for Translational Medicine and Therapeutics and the Intellectual and Developmental Disabilities Research Center at Children’s Hospital of Pennsylvania/University of Pennsylvania (National Institute of Child Health and Human Development Grant U54 HD086984. All schematics were created with BioRender.com.

## Funding

National Institutes of Health grant RO1AA027202 (S.L.B.)

National Institutes of Health grant 1F32AG079652-01A1 (N.P.)

American Parkisnon Disease Association (N.P.)

National Institutes of Health grant NIDA R37 DA053296 (J.A.D.)

National Institutes of Health grant SIG grant #S10 OD026929 (P.A)

Robert J Kleberg, Jr and Helen C Kleberg Foundation (N.B. and S.L.B.)

## Author contributions

N.P., N.M.B., and S.L.B. conceived the project. N.P. designed AAV.ACSS2 construct, performed in vitro viral transduction, hAD-tau treatment, RNA-seq, Cut&Run, DNA methylation, histone PTM mass spec, mouse injections, mouse tissue processing, WB, IHC, and IF image analysis, supervised behavioral experiments and most of the analyses. G.D., performed mouse RNA-seq, in vitro RNA-seq, and Cut&Run analysis (Fig. 2). S.Z., performed in vitro RNA-seq analysis (Fig. 1 and 3). X.L., performed DNA methylation analysis. M.G. performed Mass spec analysis. O.S. performed Drosophila experiments. N.P., designed most experiments, K.Y., and J.A.D. designed in vivo recording experiments. O.S., and N.M.B., designed drosophila experiments. H.X. provided technical advice on in vitro hAD-tau experiments. N.P., H.X., and K.Y. developed experimental protocols. C.H. performed WB, IF staining, and imaging. K.H. L.C. assisted with IHC image staining and analysis. P.A., J.A.D., V.M.L., N.M.B., and S.L.B., contributed to methodology and resources. N.P., N.M.B., and S.L.B. wrote the manuscript. All authors read and approved the paper.

## Data and materials availability

Mass spectrometry data have been deposited in the ProteomeXchange accession no. PXD048769. RNA-sequencing (GSE254996) and Cut&Run (GSE254820) data have been deposited in the National Center for Biotechnology Information Gene Expression Omnibus repository. The programming code used for data analysis is available upon request. Reagents and animal models used in this study are available upon request and execution of institutional material transfer agreement.

## Supplementary Methods

### Immunocytochemistry and high content imaging in primary hippocampal neurons

CD1 mouse hippocampi were dissected at embryonic days 16-18 and dissociated with papain (Worthington Biochemical Corporation). Neurons were resuspended in neural basal medium (Gibco, 21103) with 2% B27 (Gibco), 1x Glutamax (Gibco), and 1x Penicillin/Streptomycin (Gibco). Plates or coverslips were coated with poly-D-lysine (0.1 mg/ml, Sigma-Aldrich) in borate buffer (0.05 M boric acid, pH 8.5) overnight at room temperature. Cells were plated at a density of 50,000 cells/cm^2 surface area for all types of plates. During plating, 5% FBS was added to the cell suspension. The plating medium was replaced with neural basal medium without FBS at day 1 in vitro (DIV1). At DIV7, the plating medium was replaced with conditioned medium containing a 1:1 ratio of old and fresh medium. hAD-Tau fibrils were diluted in PBS to the desired concentrations, added on top of the cells, and incubated for 14 days until DIV 21. 1 μg tau/10^6 cells were used for AD-tau treatment. At DIV 21, cells were washed one time with PBS, extracted with 1% hexadecyltrimethylammonium bromide (HDTA, Sigma-Aldrich) for 10 min, and fixed for 10 minutes with 4% paraformaldehyde and sucrose in PBS. Fixed cells were stained with primary antibodies overnight at 4 °C and Alexa Fluor-conjugated secondary antibodies (Thermo Fisher Scientific) for 2 hrs at room temperature. Photomicrographs were obtained with an InCell scanner (GE) and quantified using the InCell analyzer software. Pooled human AD-tau was prepared from AD brain tissue as previously described (*1*) and was a gift from Virginia Lee’s lab. Post-mortem tissues were pathologically diagnosed with AD and were screened to rule out other types of comorbidities.

### Primary mouse hippocampal neurons viral treatment

For all in vitro experiments, 1.5×106 MOI of PHP.eB EF1a-ACSS2-Flag-PTV1-2A-mPlum and EF1a-mPlum were added on 5 DIV to primary neurons and cells underwent complete media change 24 hours after viral treatment and replaced with conditioned media. PHP.eB viral vectors and packaging were generated with the University of Pennsylvania (UPenn) Vector Core

### Immunocytochemistry and high-content imaging

Image analysis was performed in a blinded manner and using automated software. 21DIV hippocampal neurons were fixed with prechilled methanol at −20°C for 15 min or with ice-cold 4% paraformaldehyde (PFA) containing 1% Triton X-100 at room temperature for 15 min to remove soluble tau, as previously described. Normal tau staining was performed on neurons fixed with 4% PFA and permeabilized with 0.1% Triton X-100. After blocking with 3% BSA and 3% FBS for at least 1 h at room temperature, cells were incubated with AT8 antibody (mouse monoclonal anti-tau p-S202/T205, Thermo Fisher cat # MN1020) overnight at 4°C followed by staining with appropriate Alexa Fluor 488–conjugated secondary antibodies (Invitrogen) for 2 h at room temperature. Coverslips were mounted using Fluoromount-G containing DAPI (SouthernBiotech) to label cell nuclei. Immunofluorescence images were acquired using a microscope (BX 51; Olympus) equipped with a digital camera (DP71) and DP manager (Olympus). For the quantification of the tau pathology shown in Fig. 3F, whole coverslips were scanned using a Lamina Multilabel Slide scanner (PerkinElmer) and quantified using the image analysis platform HALO (Indica Laboratories).

### Immunofluorescence staining and imaging

All steps of immunofluorescence were carried out at room temperature unless otherwise noted. Primary hippocampal cultures were fixed in 4% PFA for 10 minutes then rinsed in DPBS 3 times for 5 min. Slides were permeabilized in 0.5% Triton x-100 and 0.01% Tween-20 in DPBS for 10 minutes and then blocked in 5% bovine serum albumin (BSA) in DPBS for 1 hour. Primary antibodies for ACSS2 (1:200; Thermo Fisher Cat. MA5-14810), NeuN (1:1000; Millipore Sigma Cat. ABN91), Synapsin (1:500, Thermo Fisher A-6442) were applied in 5% BSA in DPBS overnight at 4°C. Slides were rinsed in DPBS 3 times for 5 minutes. Secondary antibodies, Donkey anti-Rabbit IgG Alexa Flour™ 647 and Goat anti-Chicken IgY AlexaFlour™ 555 (1:500; Invitrogen Cat. A-31573 & A-21437) were applied in 5% BSA in DPBS for 1 hour. Nuclei were stained with DAPI (1:5000; Invitrogen Cat. D1306) for 10 minutes before a final wash in DPBS 3 times for 5 minutes. Slides were mounted to coverslips using 0.5% N-propyl gallate and 90% glycerol in 20 mM Tris, pH 8.0. Images were acquired using a Nikon Eclipse Ti2 Inverted Fluorescence Microscope. The images were taken under oil immersion 60x magnification (Nikon Cat# MXA22168). Quantification was performed on random-field images using ImageJ (National Institutes of Health).

### RNA-sequencing (Primary hippocampal neurons)

RNA-sequencing was performed on 9DIV and 21 DIV primary hippocampal neurons. Cells were washed 1x with PBS before RNA isolation. Total RNA was extracted using Trizol (15596018, Thermo Fisher Scientific)-chloroform following RNA purification using RNeasy mini-kit, Qiagen (QIAGEN #74106). DNA was digestion with DNase. Poly(A) + RNA was then isolated using double selection with poly-dT bead (E7490, NEB), and RNA-seq libraries were prepared using an NEB-Next Poly(A) mRNA Magnetic Isolation Illumina (E7760, NEB). Library sizes were determined on a Bioanalyzer, and concentrations were determined using the NEBNext Lib Quant Kit (E7630, NEB). Libraries were sequenced on an Illumina NextSeq 550, using paired-end sequencing of 42 bases per read.

### RNA-sequencing data analysis (Primary hippocampal neurons)

Raw paired-end reads were aligned to mm10 and refGene annotation from the UCSC Genome Browser (https://hgdownload.soe.ucsc.edu/goldenPath/mm10/) using STAR (v2.7.1a) (*2*) with parameters: --outFilterMismatchNmax 3 --outFilterMultimapNmax 1 --alignSJoverhangMin 8. Then, gene read counts were calculated using featureCounts (subread v2.0.3) (*3*) with parameters: -t exon -g gene_id -p --countReadPairs -B -Q 20. Next, we estimated unwanted variation from our dataset using RUVr function of RUVseq (v1.34.0) (*4*). Finally, differentially expressed genes were analyzed using DESeq2 (v1.40.2) (*5*). Gene ontology and Pathway analysis were performed using Enricher.

### ROSAMAP

To analyze the relevance of the AD-tau mouse primary hippocampal neuron model to human disease, differentially expressed genes in AD-tau treatment were contrasted with human AD patient RNA-seq data downloaded from AMP-AD (data sets: ROSMAP and Mayo Clinic)(*6*). Human orthologs to the 118 up-regulated and 181 down-regulated genes under AD-tau treatment were first identified using the Mouse Genome Informatics database. Each gene was evaluated for a log2(AD/healthy) patient score using average patient cohort values. The Mayo Clinic provides meta-data with calls of “AD” or “Control” for each patient, which were used to classify patients into cohorts. For the ROSMAP data, 54 patients with Braak stage 5 or 6, CERAD score 1, and cogdx score 4 were considered to be AD patients; while 27 patients with Braak stage 0, 1, or 2, CERAD score 4, and cogdx score 1 were considered to be healthy. Human patient log2(AD/healthy) fold changes were then plotted as box-and-whiskers for each AD-tau gene group and the difference between up- and down-regulated genes was evaluated statistically by Mann-Whitney.

### Cut&Run (Primary hippocampal neurons)

Cut&Run was performed similarly to previously described (*6*) on 21 DIV primary hippocampal neurons treated with hAD-tau or PBS for 14 days. Cells were detached with trypsin and collected in DMEM/F12 + 10%FBS+1% Pen/strep following centrifugation at 450xg for 3 min. Cells were counted using trypan blue to ensure >80% viability. For each condition, ∼400k cells were used. Cells were resuspended in nuclear isolation buffer (10mM HEPES-KOH, pH 7.9, 10mM KCl, 0.1% NP40, 0.5mM spermidine, 1x Halt protease inhibitor cocktail) and bound to concanavalin A lectin beads which had been washed in binding buffer (20mM HEPES-KOH, pH 7.9, 10mM KCl, 1mM CaCl2, 1mMnCl2). After bead binding, samples were split into two tubes, one for each antibody, where the binding buffer was replaced with 1:100 antibody diluted in blocking buffer (20mM HEPES-KOH, pH 7.5, 150mM NaCl, 0.1% BSA, 0.5mM spermidine, 1x Halt protease inhibitor cocktail, 2mM EDTA). H3K27ac (Active Motif 1:100. #39133), and IgG (Abcam 1:100) were added and incubated overnight at 4°C. Samples were washed in washing buffer (20mM HEPES-KOH, pH 7.5, 150mM NaCl, 0.1% BSA, 0.5mM spermidine, 1x Halt protease inhibitor cocktail), and treated with pA-MNase and CaCl2 for 30 minutes on an ice-cold pre-chilled metal block on ice. The reaction was stopped by the addition of STOP buffer (200mM NaCl, 20mM EDTA, 4mM EGTA, 50μg/mL RNase A, 40μg/mL glycogen) while gently vortexing, and DNA fragments were released by incubation at 37°C for 10 minutes. The DNA fragments were collected from bead supernatants, treated with Proteinase K for 10 minutes at 70°C, then subjected to PCI-chloroform purification. Libraries from DNA fragments were made using the NEBNext Ultra II DNA Library Prep Kit for Illumina. Library sizes were determined on a Bioanalyzer, and concentration was determined using NEBNext Lib Quant Kit (E7630, NEB). Libraries were sequenced with an Illumina NextSeq550 with 42bp per read (total of 84bp) using the NextSeq 500/550 High Output 75-cycle v2.5 kit.

### Cut&Run data analysis (Primary hippocampal neurons)

Unaligned paired-end CUT&RUN data were pooled across NextSeq lanes, then aligned using bowtie2 v2.3.4.1 (command-line parameters --local -X 1000). Data were then filtered for poor alignments using samtools v1.1 (samtools view -h -bS -f 2 -F 256 -q 5) and sorted by tag position (samtools sort -n) then PCR de-duplicated using PICARD MarkDuplicates v2.21.3-SNAPSHOT (command-line parameters REMOVE_DUPLICATES=True ASSUME_SORT_ORDER=queryname). Resulting aligned tags were processed for differential peaks using RGT-THOR v0.13.2, contrasting AD Tau-injected cells with PBS-injected cells and using matching IgG samples as background control. The FDR was controlled at 10E-10, producing 9,649 differentially enriched peaks. To make heatmaps, these were measured for RPM-adjusted tag densities in all samples and H3K27ac-IgG differences were taken, then standardized by z-score across all samples for each peak. Data were then clustered using gplots’ heatmap.2 in R. Peaks were annotated for nearest genes using HOMER v4.6 annotatePeaks.pl and this association was used to produce the boxplot comparing H3K27ac changes to gene expression changes.

### Mice

For the aging experiments, aged (21-22 months old) C57BL/6J mice were obtained from the National Institute of Aging aged rodent colonies. 10-months old C57BL/6J mice (strain # 000664), P301S transgenic mice (strain # 008169), and B6C3F1/J non-transgenic mice were procured from the Jackson Laboratory. Mice were housed in a pathogen-free barrier facility, maintained at an ambient temperature of 21-23oC and 40-50% humidity, following a 12-h light/dark cycle (from 7 am to 7 pm). Mice had ad libitum access to food and water and were accommodated in cages with no more than 4 animals each. Behavioral testing was conducted at the specified ages, followed by euthanasia cervical dislocation. All experimental procedures involving animals were approved by the Institutional Animal Care and Use Committee (IACUC) under protocols 804849 and 804484. All personnel involved in the experiments have received appropriate training and qualifications in accordance with the guidelines of the Animal Welfare Act (AWA) and the Public Health Service (PHS) policy.

### Retro-orbital injection

C57BL/6J mice were injected via retro-orbital injection of PHP.eB 2.0 x1011 viral genome (vg) of EF1a-ACSS2-Flag-PTV1-2A-mPlum (AAV.ACSS2) and EF1a-mPlum (vector) constructs. These AAVs were produced by Penn Vector Core.

### Stereotaxic injection

P301S mice aged 2.5 months were anesthetized by isoflurane gas and securely positioned within a sterile field within a stereotaxic apparatus. To ensure proper lubrication, artificial tears were applied to the eyes. The skin was disinfected with betadine solution and a small incision was made to expose the skull. A Hamilton syringe loaded with the viral vector was carefully inserted into the dorsal hippocampus (AP, −2.0mm; DV, −1.4mm; ML, +/−1.5mm from Bregma). Each mouse was bilaterally injected (2 ul per injection site) with a total of 1.50×1010 gc of PHP.eB-EF1a-ACSS2-Flag-PTV1-2A-mPlum (AAV.ACSS2) and PHP.eB-EF1a-mPlum (vector).

### Behavioral Assay

All behavioral assays were performed in a blinded manner, and the individual collecting the data was unaware of both group assignments and received interventions. All behavioral experiments were conducted between 7 am and 11 am to minimize potential time-of-day effects.

### Fear Conditioning

Mice were handled for three consecutive days for 2 minutes each before the start of the experiment. On the day of fear acquisition, mice were individually placed in conditioning chambers (Med Associates). Aged C57BL/6J mice (21-22 months old) were subjected to 24-hour cued, 48-hour contextual, 2 weeks (remote) cued, and contextual recall. Mice were habituated to the novel environment for 2 minutes. Auditory cue (an un-modulating tone: 85 dB and white noise) was presented for 30 seconds, co-terminating with a mild 2s, 1.0 mA foot shock, and repeated for a total of 3 times. Chambers were wiped down with 70% ethanol between each round (for a schematic, see Fig. 6I). Mice were promptly removed 1 minute after the last shock and returned to their home cages. To assess cued recall, chambers were modified to remove spatial and olfactory cues. The shape of the chamber was modified with cardboard inserts, and the barred flooring was replaced with a solid layer. The scent of ethanol was masked with vanilla extract. The same tone (minus the shock) was repeated in the same pattern as the previous day. Freezing behavior was monitored automatically using FreezeScanTM software (CleverSys, Inc.) for the entire recall period. The 10-month-old C57BL/6J mice underwent the same fear condition paradigm as aged mice but without recall. These mice were euthanized 30 minutes post acquisition. The P301S mice undergoing the contextual fear conditioning paradigm, mice were habituated to the chamber for 3 minutes. 3x mild 2s, 0.8 mA foot shock (for a schematic, see Fig. 4, A and B) with 80dB background noise. 24 hr later, the freezing response was tested for 5 consecutive minutes with 80dB background noise. Mice were euthanized 30 minutes post context recall.

### Y-maze test of spontaneous alternation

Aged C57BL6/JC mice (21-22 months) were placed at one end of a Y-shaped arena and allowed to explore freely for 6 minutes. The scoring of arm entries was conducted visually. An arm entry was counted when the hindquarters of the mouse fully entered the new arm. The percentage of spontaneous alternations was computed using the following formula: SAR = [Number of alternations/(total arm entries – 2)] x 100.

### Open Field

P301S mice were introduced into an open arena measuring 30 cm x 40 cm and were allowed to explore freely for 6 minutes. Videos of their behavior were subsequently subjected to analysis utilizing the ANY-mazeTM behavioral tracking software (Stoelting Co.). To quantify locomotion, the total path length covered within the entire 6-minute interval was assessed in meters. Thigmotaxis, which reflects the tendency of the mice to stay close to the periphery, was quantified by calculating the ratio of time spent within the peripheral zone (8 cm distance from the arena walls) to the total elapsed time.

### Immunohistochemistry mouse tissue and image analysis

Mouse brains were perfused with PBS at a flow rate of 2 mL/min for 15 min and immersion-fixed with 4% paraformaldehyde for 6 hours at 4C. The tissue was transferred to 30% sucrose in DPBS overnight at 4C. The tissue was briefly rinsed in DPBS before being embedded in OCT (Fisher Sci Cat. 23-730-571). The tissue was stored at −80C until processed. Sections of 6-μm thickness were cut and used for immunostaining. After deparaffinization in xylene and a decreasing series of ethanol dilutions, the tissue was rinsed in water and incubated in 3% H2O2 at room temperature for 10 minutes to block endogenous peroxidase activity. Following a 1xPBS wash, the tissue is blocked in horse serum/BSA blocking buffer for 1h at room temperature. Primary antibodies were diluted in blocking buffer and sections were incubated overnight at 4oC (AT8 (Mouse monoclonal anti-tau p-S202/T205, Thermo Fisher cat # MN1020, 1:1000), NeuN (A60) (for neuronal count, Millipore cat # MAB377, 1:1000) or ACSS2 (Thermo Fisher, cat # MA5-145810, 1:200). The next day, tissue is washed in 1xPBS and incubated in secondary antibody conjugated with HRP (Vector Labs) for 2-hour. Following 3x 1xPBS washes, the staining is developed using DAB reagents (Vector Labs). Finally, the tissue is counterstained with hematoxylin and re-hydrated through increasing ethanol series and xylene before being mounted with Permount. Quantification was performed within the region of interest (ROI) using a threshold-based classifier by QuPath software. All the conditions to be compared were quantified using the same classifier. For IHC, 3-4 sections were quantified for the hippocampus. The NeuN quantification was normalized to the overexpression group. The average of the overexpression group was set to 100% to remove staining variation at different slides.

### LTP and in vivo recording

Mice were deeply anesthetized with ketamine/xylazine, and then the mice were transcardially perfused with ice-cold, well-oxygenated (95% O2/5%CO2) N-methyl-D-glucamine (NMDG) based artificial cerebrospinal fluid (ACSF, in mM): 92 NMDG, 2.5 KCl, 1.2 NaH2PO4, 30 NaHCO3, 20 HEPES, 25 glucose, 2 thiourea, 5 Na-ascorbate, 3 Na-pyruvate, 0.5 CaCl2, and 10 MgSO4, pH 7.3-7.4 with concentrated HCl (*7*). Sagittal brain slices containing the dorsal hippocampus (200 µm) were cut in ice-cold, well-oxygenated (95% O2/5%CO2) NMDG ACSF with a Leica VT1200S vibratome. The slices were allowed to recover for 13 min. in 32°C NMDG. Slices were then kept in a HEPES-based holding ACSF solution (in mM): 92 NaCl, 2.5 KCl, 1.2 NaH2PO4, 30 NaHCO3, 20 HEPES, 25 glucose, 2 thiourea, 5 Na-ascorbate, 3 Na-pyruvate, 2 CaCl2, and 2 MgSO4 at room temperature for at least 1 hour before beginning to record. The dorsal hippocampal slices were continuously bathed at 32–34 °C in well-oxygenated standard recording ACSF (in mM): 124 NaCl, 2.5 KCl, 1.2 NaH2PO4, 24 NaHCO3, 5 HEPES, 12.5 glucose, 2 CaCl2, and 2 MgSO4. CA1 pyramidal neurons were recorded with 2-3 MΩ electrodes filled with potassium-based solution (in mM): 135 K-methanesulfonate, 4 NaCl, 10 HEPES, 0.06 EGTA, 0.01 CaCl2, 2 MgCl2, 2 Na2-ATP, 0.3 Na-GTP, and 0.15 spermine pH 7.3 (*8*). All recordings were made in the presence of 100 µM picrotoxin to inhibit GABA activity. For most of the recordings, 5-10 min of tight-seal, cell-attached recording was maintained to observe spontaneous action potential firings before a whole-cell configuration was achieved by applying a brief suction. We obtained an input-output relationship first to determine the stimulation intensity to study synaptic plasticity. In general, the baseline EPSP amplitude was at 20% of the generated input-output curve. EPSPs were evoked using 0.1 ms current pulses every 30 s with a tungsten bipolar stimulating electrode (WPI, Sarasota, FL) placed in stratum radiatum to stimulate the Schaffer collateral pathway. To induce long-term synaptic potentiation (LTP), 3 trains of high-frequency stimulation (*8*) (HFS, 100 Hz for 1s with 20 s intervals) were applied immediately after recording a stable 5 min baseline. The LTP induction protocol was limited to once per slice. All slices were cut between CA1 and CA3 to prevent the propagation of recurrent epileptiform activity.

### Western blot

Cells and tissue were lysed in RIPA buffer containing 50 mM Tris pH 8.0, 0.5 mM EDTA, 150 mM NaCl, 1% NP40, 1% SDS, supplemented with HALT protease and phosphatase inhibitor cocktail (Life Technologies, number 78446) and sodium butyrate (10 mM). Protein concentration was determined by Bradford protein assay, and equal amounts of protein were loaded onto 4–12% Bis-Tris polyacrylamide gels (NuPAGE). Proteins were transferred to PVDF membrane and subsequently blocked with 5% milk in TBS-T. Membranes were incubated with primary antibodies ACSS2 (Cell Signaling, Cat # 3658 1:1000), Actin (Cell Signaling Cat#4970 1:2000), H3K27ac (Abcam ab4729, 1:1000), H3K9ac (Active Motif, RRID AB_2561017 1:1000) diluted in blocking buffer for 4 °C overnight. Membranes were washed three times in TBS-T for 10 minutes each before incubation with HRP-conjugated secondary antibodies in blocking buffer. Membranes were washed again as before, developed with SuperSignal West pico PLUS chemiluminescent substrate (Thermo Fisher) then imaged with an Amersham Imager 600.

### RNA-sequencing (Mouse)

Flash frozen brains were dissected to isolate whole hippocampi (Fig. 6D) or 1 mm-thick slices of the dorsal hippocampus (Fig. 6A and Fig. 5A). Tissue was homogenized in TRIzol reagent using a dounce homogenizer. Following homogenization, chloroform was introduced to the samples, which were then subjected to centrifugation at 14,000g for 15 minutes at 4 °C. Next, RNA was isolated according to the manufacturer’s protocol (RNeasy mini-kit, Qiagen). Total RNA quality was assessed on the Bioanalyzer platform using the RNA 6000 Nano assay (Agilent). Poly(A) + RNA was then isolated using double selection with poly-dT bead (E7490, NEB), and RNA-seq libraries were prepared using an NEB-Next Poly(A) mRNA Magnetic Isolation Illumina (E7760, NEB). Library sizes were determined on a Bioanalyzer, and concentrations were determined using the NEBNext Lib Quant Kit (E7630, NEB). Libraries were sequenced on an Illumina NextSeq 550, using paired-end sequencing of 42 bases per read.

### RNA-sequencing data analysis (Mouse)

RNA-seq data were prepared for analysis as follows: NextSeq sequencing data was demultiplexed using native applications on BaseSpace. Demultiplexed FASTQs were aligned by RNA-STAR 2.5.2 to assembly mm10 (GRCm38). Aligned reads were mapped to genomic features using HTSeq 0.9.1. Quantification, and library size adjustment, were performed using DESeq2. RUVseq was utilized to remove unwanted variation. Data were first filtered for low-expressed genes, keeping those with more than 5 tags in at least two samples, then subjected to upper quartile normalization, then regressed to the normalized subtracted cue recall (aged mice) and context recall (PS19 mice) behavioral scores. Residuals were calculated using edgeR after estimating common and tag-wise dispersion and then these were used as input to RUVr, calibrated to calculate the per-sample coefficients for a single source of unwanted variation. The resulting coefficients were then used in a design ∼unwanted+behavior with DESeq2 to obtain differentially expressed genes. The significance of gene alterations was determined using the Wald test with multiple test corrections according to the BenjaminiHochberg method with FDR < 0.1. Gene ontology and pathway analyses were performed using the Enricher (*9, 10*). Figures generated using R, Tableau, Prism, and SRPlot (*11*).

### Histone post-translational modification mass spectrometry

Histone extraction from frozen mouse hippocampi was performed as previously described (*12*). Procedures were performed on ice or at 4 °C. The nuclei were isolated by suspending sample type into nuclei isolation buffer (15mM Tris–HCl (pH 7.5), 15mM NaCl, 60mM KCl, 5mM MgCl2, 1mM CaCl2, 250mM sucrose, 0.3% NP-40) including the following inhibitors: 1mM DTT, 0.5mM AEBSF and 10mM sodium butyrate. Nuclei were separated by centrifugation (600 g for 5 min) and 50 µL of cold 0.4N H2SO4 was added to the nuclei pellet. Nuclei were incubated at 4°C with shaking for 3 hours without rotation, pelleted at 14000 g for 10 min, and histones were precipitated from the supernatant with 33% TCA (w/v). Purified histones were resuspended in 20ul of 50mM NH4HCO3, pH 8.0. Derivitization and digestion were performed using the bottom-up strategy described previously (*12*). Briefly, derivatization of the lysine residues side chains was performed by preparing a mix of pure propionic anhydride: acetonitrile (1:3); 15ul of this mix was added to the histone sample, followed by immediate addition of 3ul of ammonium hydroxide to adjust the pH to 8.0 after the introduction of propionic anhydride to the mixture. The reaction was performed twice. Digestion was performed using trypsin at an enzyme: sample ratio of 1:20, overnight at room temperature. The derivatization reaction was repeated to derivatize peptide N-termini. Samples were then desalted using Pierce C18 Tips, 10 µL bed (Thermo Scientific).

### Bottom-up nanoLC-MS/MS and data analysis

Histone tail peptides were analyzed in 3 biological replicates with nanoLC-MS/MS. Peptides were separated using an UltiMate3000 (Dionex) HPLC system (Thermo Fisher Scientific, San Jose, Ca, USA) using a 75 μm ID fused capillary pulled in-house and packed with 2.4 μm ReproSil-Pur C18 beads to 20 cm. The HPLC gradient was 0%–34% solvent B (A = 0.1% formic acid, B = 95% acetonitrile, 0.1% formic acid) over 46 min and from 34% to 90% solvent B in 5 min at a flow rate of 300nl/min. The QExactive HF (Thermo Fisher Scientific, San Jose, CA, USA) mass spectrometer was configured following the data-independent acquisition (DIA) method previously described (*12*). Full scan MS spectra (m/z 300-1100) were acquired with a resolution of 60,000 with an AGC target of 1e6; MS/MS spectra were acquired with 50 m/z precursor isolation windows (stepped CID normalized collision energy of 25, 27.5, 30, and an AGC target of 5e5. Mass spectrometry data files were imported into Skyline to calculate integrated MS2 peak areas. For statistical analysis, a 2-tailed t-test was performed (significant if p < 0.05).

### DNA extraction and DNA methylation sequencing

Frozen tissue was homogenized using SDS buffer (0.1M Tris, pH = 8, 0.2M NaCl, 5mM EDTA pH = 8, 0.2% SDS, and 0.2mg/ml Proteinase K added just before use) and digested for 1 hour at 55 C water bath. DNA extraction was performed performed using Phenol:chloroform: isoamyl alcohol solution (PCI) (25:24:1, v/v/v) (Invitrogen, Cat # 15593049). An equal volume of PCI was added to the sample and vortexed for 1 min. Samples were centrifuged at RT for 5 min at 16,000xg. The upper aqueous phase was transferred to a fresh tube. PCI was added for the second time to obtain pure extract. After the final centrifugation, DNA was precipitated using 100% Ethanol, 7.5 M NH4OAC, and 20 mg/ul Glycoblue. Samples were placed in −80 overnight and centrifuged at 4C for 1 hour and max speed to pellet the DNA. The pellet was 2x washed with 70% ethanol and air dried at RT for 5 min. The DNA pellet was resuspended in 10 mM Tris-HCl, pH 8.5, and quantified using Qubit. DNA methylation was sequenced using Illumina Infinium Mouse Methylation BeadChip Kit (20041558, Illumina).

### DNA methylation data analysis

DNA methylation array data was analyzed by Sesame (*13*) V1.18.4. Data was preprocessed with Sesame preparation code “TQCDPB” designed for mouse MM285 array. Beta value was generated for visualization like heatmaps and PCA. Differential methylation locus (DML) and differential methylation regions (DMR) were obtained using Sesame function DML and DMR. P-values were further adjusted to false discovery rate (FDR). PCA was generated using scikit-learn V 0.23.1. (*14*). Probes were mapped to mm10. CpGs/regions were annotated to genomic features using R package ChIPpeakAnno (*15*) V 3.34.1 and annotated to genes using homer v4.11129. GO analysis is done using shinyGO 0.77 (http://bioinformatics.sdstate.edu/go/).

### Fly husbandry and fly lines (Drosophila)

Flies were maintained on a standard yeast/cornmeal-based medium at 25°C on a 12hr:12hr light–dark cycle. Elav-Gal4 (#458), UAS-dACSS2-RNAi (#41917), UAS-mCherry-RNAi (#35785), UAS-GFP (#5137) lines were obtained from the Bloomington Drosophila Stock Center. The pUAST-attB-dACSS2-HA construct was used to make the transgenic fly line (BestGene). For lifespan analysis, adult male flies were collected at eclosion and were flipped to fresh food vials every 2 days. The number of dead/censored flies was recorded after flipping.

### Climbing assay

Males of the appropriate age were placed into the empty vials and allowed to acclimatize for 30 min. Vials were gently tapped 3 times to bring flies to the bottom, and a video was captured for 30 seconds after the taps. The distance in cm that flies climbed in 30 seconds was counted. The experiment was repeated 3 times for each fly.

---

**Fig S1. related to Fig 1.**
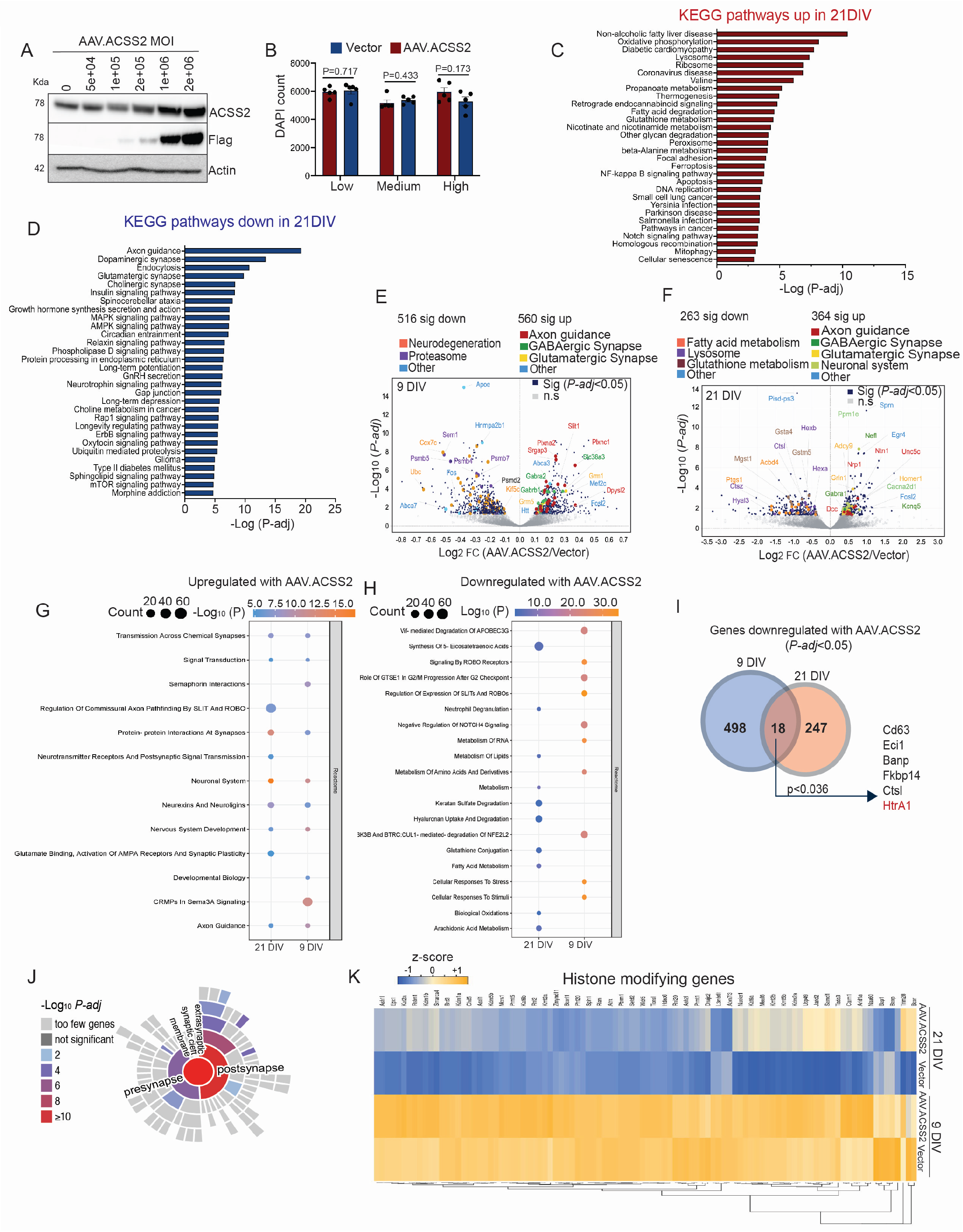
**A.** WB showing ACSS2 upregulation in a dose dependent manner 7 days post transduction, MOI (Multiplicity of infection). **B**. Percentage of hippocampal neuron survival with low (8.50E+03), medium (8.50E+04), and high (8.50E+05) MOI. **C.** Top 30 upregulated and **D**. downregulated KEGG pathways. **E.** Volcano plots from RNA-seq of 9 DIV and **F**. 21 DIV hippocampal neurons. Dark blue dots are significant genes (*P-adj* <0.05). Significant genes of interest are color-coded according to their respective Reactome pathways. **G.** Enriched Reactome pathway analysis of the list of upregulated and, **H**. downregulated DEGs (*P-adj* <0.05) from volcano plots in Fig. S1, J and K. **I**. Venn diagram showing the overlap of DEGs (*P-adj* <0.05) downregulated by AAV.ACSS2/vector in 9 and 21 DIV hippocampal neurons, permutation test *P < 0.036*. Gene list (black) ranked based on fold change; the red colored gene is AD related. **J.** Sunburst plots depict SynGo ontology terms organized by synaptic locations using the list of genes from Fig 1J. **K.** Heatmap showing the average normalized expression of 100 epigenetic regulating enzymes and chromatin modifier genes (*P-adj* <0.05) across n=3 biological replicates. The list of genes was curated based on known canonical genes.

**Fig S2. related to Fig 2.**
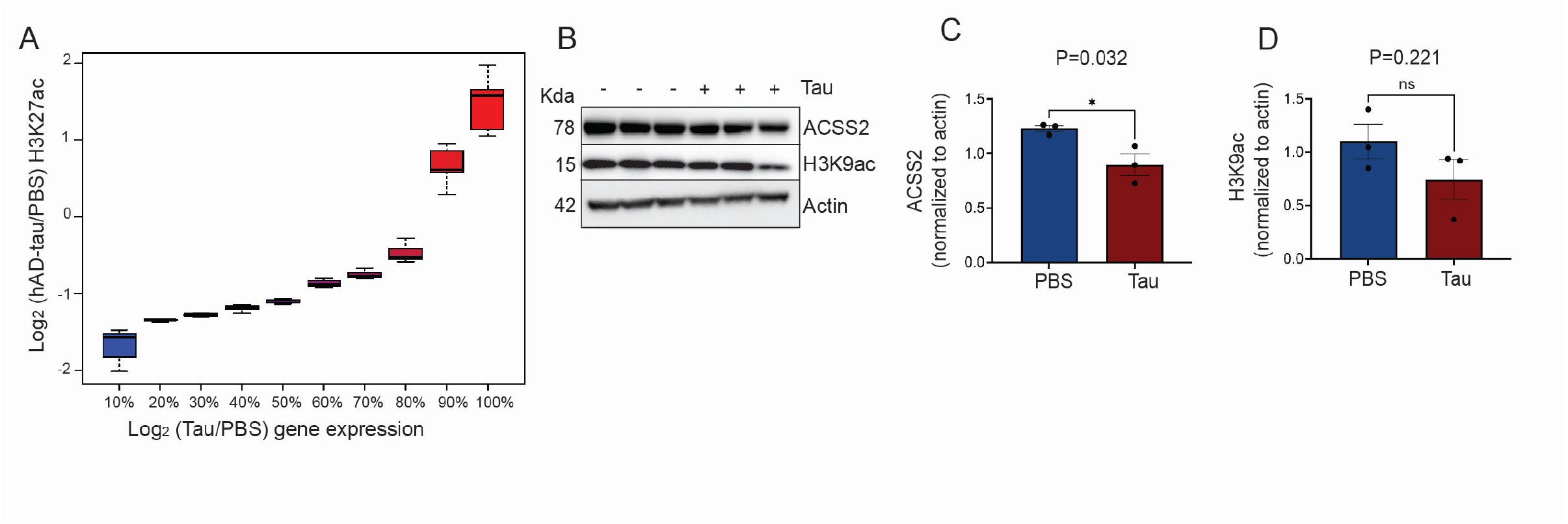
**A.** Construction plot that determines the binning of gene expression data into low and high bins related to Fig. 2H. **B**. Reduced ACSS2 and H3K9ac with AD-tau treatment in 21 DIV primary hippocampal neurons. **C**. WB quantification: ACSS2 and **d**. H3K9ac levels normalized to actin, n=3 biological replicates, mean ± s.e.m*, unpaired t-test*, P*<0.05.

**Fig S3. related to Fig 3.**
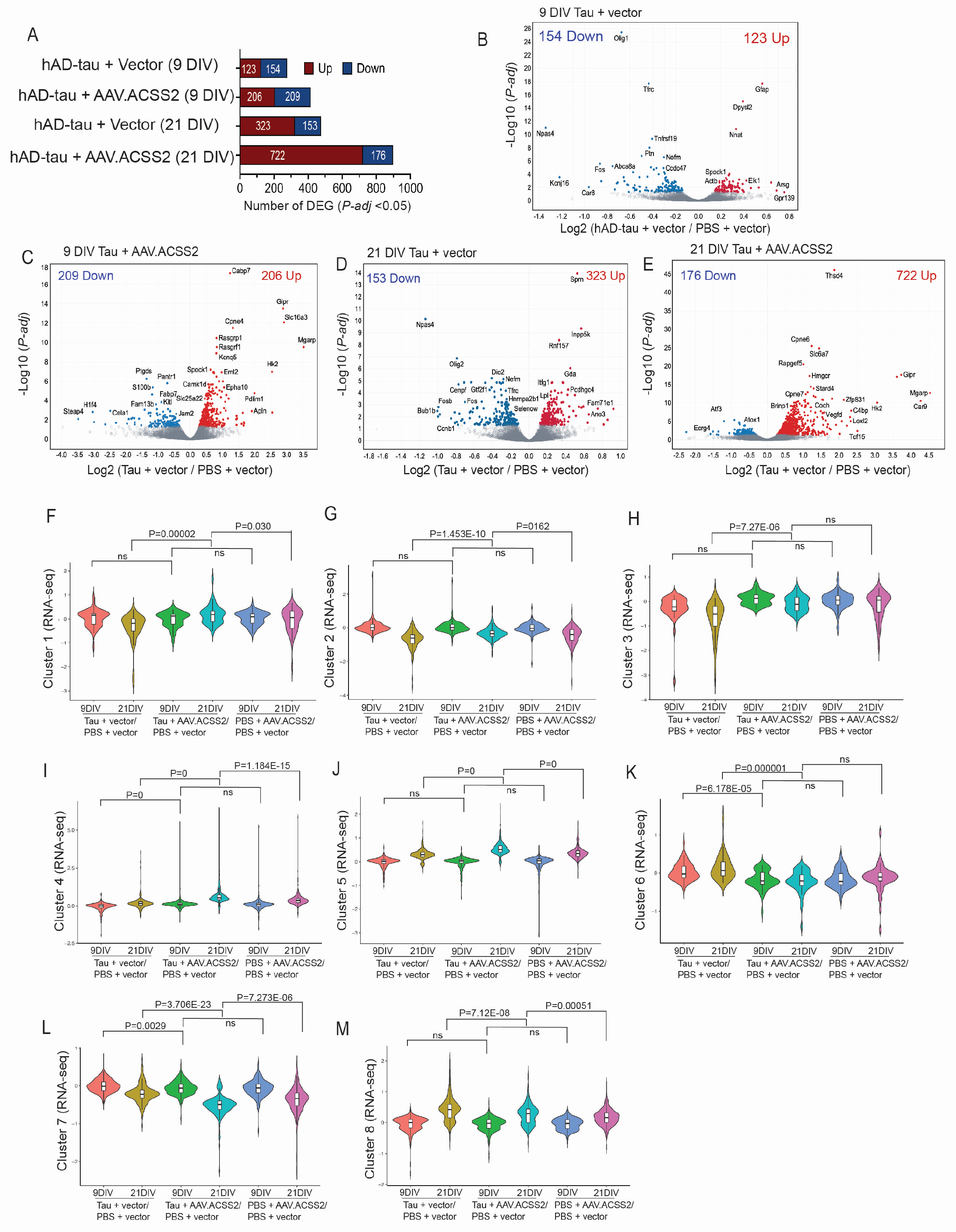
**A.** Increased transcriptional response with AAV.ACSS2 in 21 DIV hippocampal neurons treated with hAD-tau. Stacked bars show the number of DEGs (*P-adj <* 0.05) per comparison; number of DEGs significantly upregulated (red) and downregulated (blue). **B.** Volcano plot of RNA-seq data from 9 DIV hippocampal neurons (hAD-tau + vector/ PBS + vector), **C.** 9 DIV (hAD-tau + AAV.ACSS2/ PBS + vector), **D.** 21 DIV (hAD-tau + vector/ PBS + vector), **E.** 21 DIV (hAD-tau + AAV.ACSS2/ PBS + vector) showing DEGs (*P-adj* < 0.05) that are upregulated (red) or downregulated (blue). Non-significant genes are colored gray. Genes of interest are labeled. **F-K**. Violin plots of the log2(AAV.ACSS2/vector) with PBS-only for each cluster (no standardization), contrasting to the LFCs from the hAD-tau + AAV.ACSS2 columns. FDR control with Benjamini-Hochberg with the cutoff the same.

**Fig S4. related to Fig 4.**
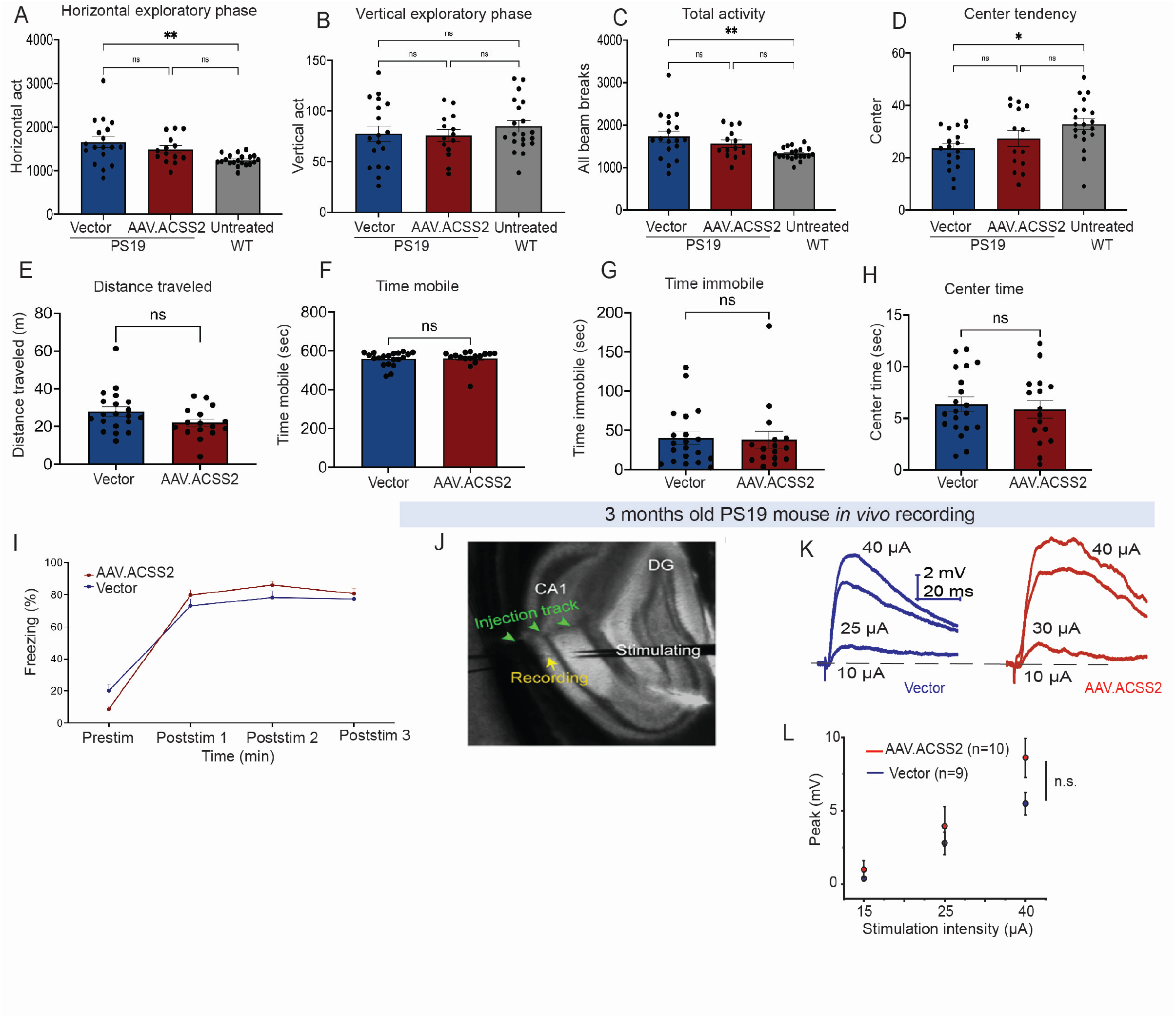
**A-H**. Open field test in PS19 mice. **A.** horizontal exploratory act, **B**. vertical exploratory act, **C**. total activity, **D.** central tendency. For A-D beam break was analyzed, n= 18 PS19 vector and n=14 PS19 AAV.ACSS2, and n= 20 untreated wildtype. *One-way ANOVA*, **P<0.01**. E.** Distance traveled, **F.** time spent mobile showing no motor deficits, **G.** time spent immobile, and **H**. time spent in the center of the field showing no difference in anxiety level. For G and H, ANY-maze was used to automatically track and quantify mouse behavior during open field test. Analysis was performed only for AAV.ACSS2, and vector treated PS19 mice. **I.** FC acquisition shows no difference between AAV.ACSS2 and vector treated PS19 mice. The same mice used for the open field were used for FC. Related to Fig. 4B and C. **J-L.** Overexpression of ACSS2 does not change the basal synaptic strength in CA1 pyramidal neurons. **J**. *In vivo* recording in mouse dorsal hippocampus. **K.** Representative traces of input/output measurements of EPSPs recorded in mice either from vector controls (left, green) or from ACCS2 overexpressing mice (right, red). **L.** Summary of the absolute EPSP peak amplitudes evoked by corresponding intensities of stimulus. Related to Fig. 4D-K.

**Fig S5. related to Fig 5.**
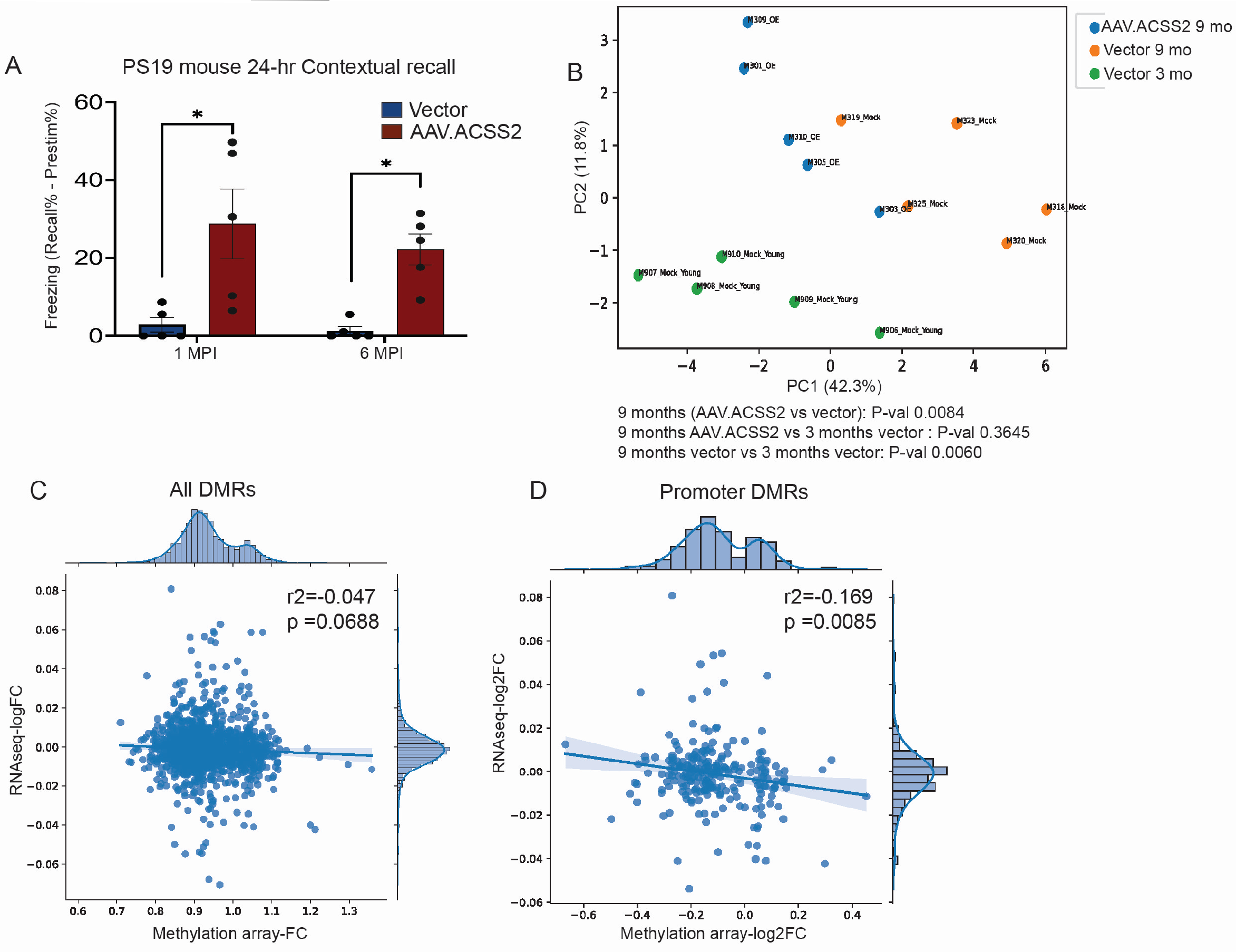
**A.** 24-hour contextual fear recall in PS19 male mice at 1 month and 6 months post stereotaxic injection, related to Fig 5A-C. **B**. PCA plot of differentially methylated regions showing a separation between aged P19 mice treated with vector (mock), and aged PS19 with AAV.ACSS2, and PS19 young with vector (mock) treatment. P value was computed using the coordinate of each sample in PC1, with two tailed Welch *t-test*.

**Fig S6. related to Fig 6.**
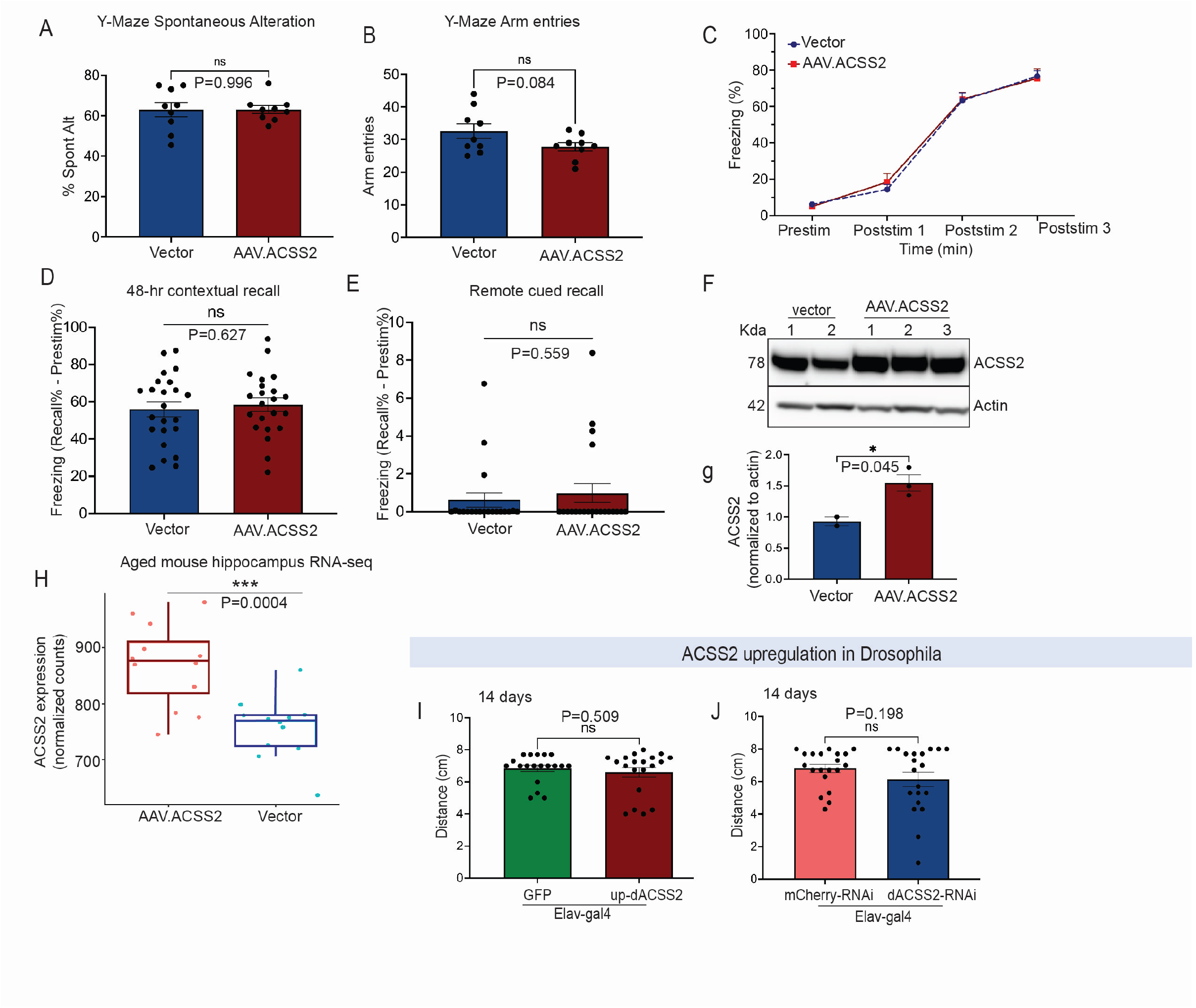
**A**. ACSS2 upregulation does not impact short-term memory. % Spontaneous alteration in aged 22-23 months old mice 2 months post systemic administration. **B**. Number of arm entries highlighting no motor deficits. **C.** FC acquisition shows no difference between AAV.ACSS2 and vector treated aged mice. **D**. 48-hour contextual recall, **E**. 2 weeks remote cued recall. **f**. WB using lysate from aged mouse prefrontal cortex, **G**. ratio of ACSS2 to actin quantified using ImageJ (n=3 AAV.ACSS2 and n=2 vector; unpaired *t* test, mean ± s.e.m, P*<0.05). **H.** ACSS2 gene expression level in AAV.ACSS2 treated aged mice from the RNA-seq data in Fig 6m. **I.** 14 day climbing assay in Drosophila with neuronal up-dACSS2, **j.** and neuronal dACSS2 knockdown.

